# Sema7A and Sema4D Heterodimerization is Essential for Membrane Targeting and Neocortical Wiring

**DOI:** 10.1101/2023.02.10.527998

**Authors:** Paraskevi Bessa, Andrew G. Newman, Kuo Yan, Theres Schaub, Rike Dannenberg, Denis Lajkó, Julia Eilenberger, Theresa Brunet, Kathrin Textoris-Taube, Emanuel Kemmler, Priyanka Banerjee, Ethiraj Ravindran, Michael Mülleder, Robert Preissner, Rudolf Grosschedl, Marta Rosário, Victor Tarabykin

**Affiliations:** Institute of Cell Biology and Neurobiology, Charité-Universitätsmedizin Berlin, corporate member of Freie Universität Berlin, Humboldt-Universität zu Berlin, and Berlin Institute of Health, Charitéplatz 1, 10117 Berlin, Germany; Dr. v. Hauner Children’s Hospital, University Hospital, Ludwig-Maximilians-University Munich, 80337 Munich, Germany; Department of Pediatric Neurology, Dr. v. Hauner Kinderklinik, University of Munich, Munich, Germany; Institute of Human Genetics, Klinikum rechts der Isar, School of Medicine, Technical University of Munich, Munich, Germany; Institute of Biochemistry, Charité -Universitätsmedizin Berlin, corporate member of Freie Universität Berlin, Humboldt-Universität zu Berlin, and Berlin Institute of Health, Philippstrasse 12, 10115, Berlin, Germany; Core Facility – High-Throughput Mass Spectrometry, Charité – Universitätsmedizin Berlin, Corporate Member of Freie Universität Berlin and Humboldt-Universität zu Berlin, Core Facility – High-Throughput Mass Spectrometry, Am Charitéplatz 1, Berlin, Germany; Institute of Physiology, Charité -Universitätsmedizin Berlin, corporate member of Freie Universität Berlin, Humboldt-Universität zu Berlin, and Berlin Institute of Health, Philippstrasse 12, 10115, Berlin, Germany; Department of Cellular and Molecular Immunology, Max Planck Institute of Immunobiology and Epigenetics, 79108 Freiburg, Germany; Research Institute of Medical Genetics, Tomsk National Research Medical Center of the Russian Academy of Sciences, Tomsk, 634009, Russia; Institute of Neuroscience, University of Nizhny Novgorod, pr. Gagarina 24, Nizhny Novgorod, Russia

**Keywords:** Axonal growth, neuronal migration, neocortex development, Semaphorin4D, Semaphorin7A

## Abstract

Disruption of neocortical circuitry and architecture in humans causes numerous neurodevelopmental disorders. Neocortical cytoarchitecture is orchestrated by various transcription factors such as Satb2 that control target genes during strict time windows. In humans, mutations of SATB2 cause SATB2 Associated Syndrome (SAS), a multisymptomatic syndrome involving intellectual disability, speech delay, epilepsy and craniofacial defects. We show that Satb2 controls neuronal migration and axonal outgrowth by inducing the expression of the GPI-anchored protein, Sema7A. We find that heterodimerization with Sema4D increases targeting of Sema4D to the membrane and is required for Sema7A function. Finally, we report that membrane localization and pos- translational modification of the Sema7A-Sema4D complex is disrupted by a novel de novo mutation in Sema4D (Q497P) that is associated with epilepsy in humans.

**GRAPHICAL ABSTRACT:** 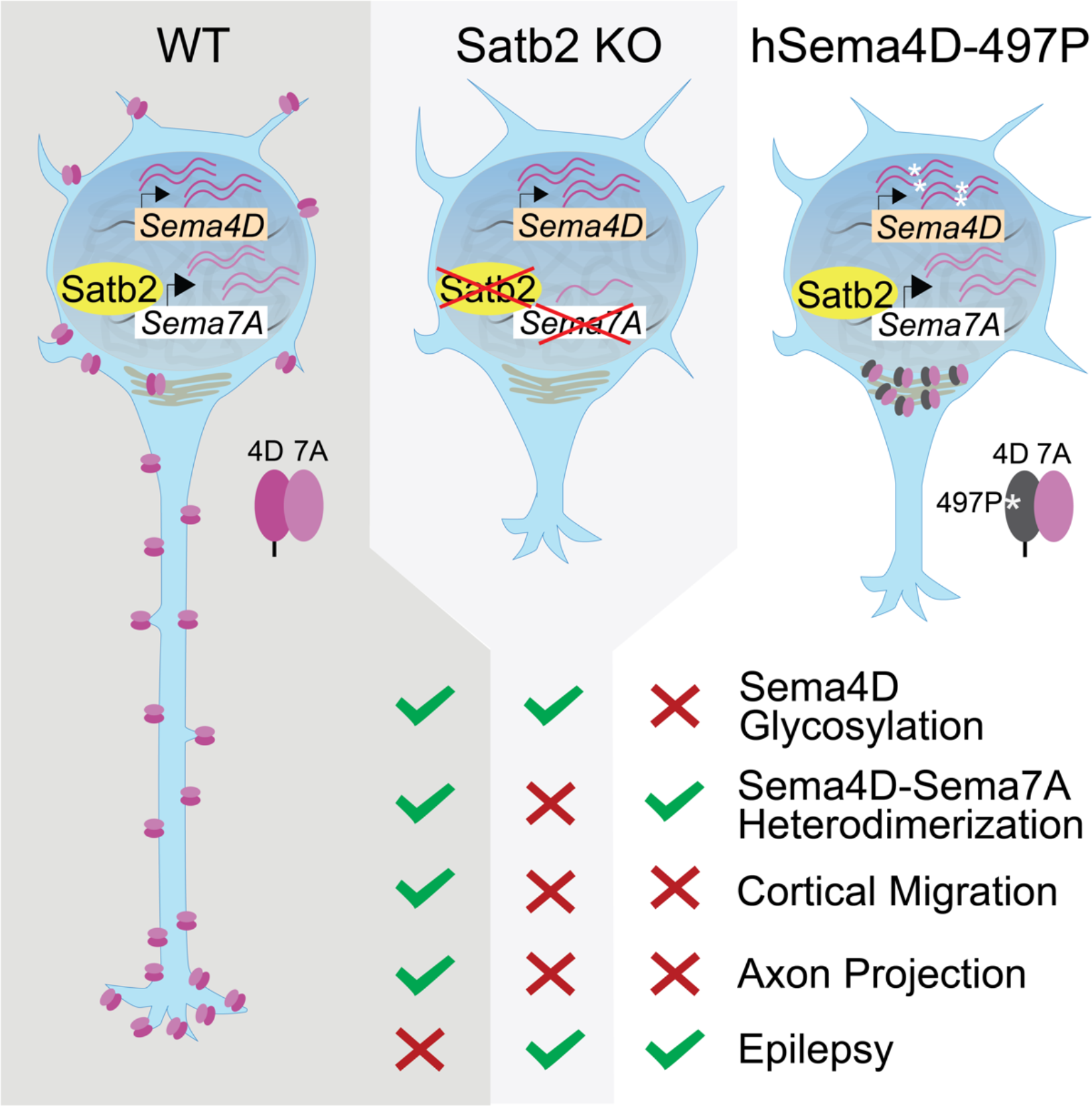

**HIGHLIGHTS:**

- Sema7A is a direct Satb2 target that drives neuronal migration and axon outgrowth
- Sema7A exerts its effect by heterodimerizing with Sema4D at neurites and growth cones
- Sema7A increases cell surface localization of Sema4D
- *De novo* human Sema4D-Q497P mutation causes epilepsy, inhibits post-translational processing & surface localization

**eTOC:** Sema7A is a direct target of the transcription factor Satb2. Sema7A promotes normal migration and axon outgrowth in cortical neurons by modulating reverse signaling via Sema4D. These processes are dependent on Sema7A-Sema4D heterodimerization and membrane localization; insufficient transcription of Sema7A or incomplete glycosylation of Sema4D inhibit this progression.

## INTRODUCTION

Accurate axonal navigation and neuronal migration are critical for the establishment of functional neocortical circuits. Both specification and initial extension of neocortical axons occurs in the subventricular zone (SVZ) and intermediate zone (IZ) of the developing neocortex in a process of neuronal polarization ^1, 2^. There are two major decisions that every neuron in the SVZ must make: first, which neurite will become an axon and which a leading process, and second, in what direction its axon will extend: medially, to join cortico-cortical axonal bundle or laterally, to join corticofugal tracts that leave the neocortex.

In the last two decades, several transcription factors have been identified, whose ablation causes abnormal development of projections of neocortical neurons (see reviews from ^3–6^). Deletion of the *Satb2 gene* (Special AT-rich sequence binding 2) causes failure of callosal neurons to form the corpus callosum ^7, 8^. In humans, mutations of SATB2 gene cause SATB2 Associated Syndrome (SAS) and characterized by symptoms such as developmental delay (DD)/ intellectual disability (ID), absent or limited speech development, epilepsy, craniofacial abnormalities including palatal and dental abnormalities, dysmorphic features and behavior ^9–12^.

It is widely known that transcription factors control long-range neocortical projection neurons to reach their target regions, by regulating the transcription of receptor-ligand pairs that can interpret environmental signals. Previously, we identified Unc5C and DCC as targets of Satb2 and Ctip2 transcription factors that control a binary choice of layer V neurons to project either medially or laterally ^13^. However, these molecules were not involved in the control of proper development of layer II-III projections that constitute most cortico-cortical connections.

The Semaphorin family of axon guidance molecules are important for the development of the nervous system as well as in immune cell function. Their fundamental role in growth cone collapse and axon fasciculation has been thoroughly studied in different model organisms ^14–16^. Mammalian Semaphorins are divided into five subfamilies consisting of secreted and membrane-bound proteins ^17–19^. Semaphorin7A (Sema7A) is the only Semaphorin that is attached to the cell membrane via a C-terminal Glyco- Phosphatidyl-Inositol (GPI) anchor and lacks a cytoplasmic domain. Semaphorin 7A forms homodimers through SEMA and IG domain interactions and dimerization seems to be important for the Semaphorin function in different cell systems ^20, 21^.

Classically, Semaphorins have been observed to act as ligands and stimulate signal transduction by binding *in trans* to their Plexin or Neuropilin receptors ^15, 22, 23^. During the past few years, evidence is mounting for the existence of reverse Semaphorin signaling, whereby ligand- binding activity induces signal transduction in the Sema-containing cell. Reverse signaling has been reported for Sema6A ^24^, Sema6B ^25^, Sema6D ^26^ and Sema4A ^27^ and have extended implications for neuronal development and axonal pathfinding ^28^. Semaphorins that can signal in reverse all have cytoplasmic domain.

Here, we identify Sema7A as a Satb2 direct target, that cell-intrinsically controls both radial migration and axon outgrowth of layer II-III neocortical neurons. Additionally, we report that Sema7A forms a heterodimer with Sema4D. Sema4D-Sema7A interaction is required for both the migration and axonal elongation of upper layer (UL) neocortical neurons. Moreover, we discovered the *de novo* mutation hSema4D- Q497P in a patient with epilepsy. This mutation interferes with the post-translational glycosylation of Sema4D and thus disrupts the localization of Sema4D:Sema7A heterodimers to the plasma membrane and growth cones. Furthermore, the hSema4D-497P mutation inhibits migration and axon projections of cortical neurons in the murine developing neocortex. Our results emphasize the importance of stabilizing/regulating the membrane localization of Sema7A and demonstrate the role of Sema4D and residue 497 in this process. Overall, we identify a crucial role of the Sema4D-Sema7A heterodimer in initiating neuronal migration and axonal growth downstream of Satb2. Our data shed light on new mechanisms where traditionally ligand-considered molecules, such as Sema4D and Sema7A, form signaling complexes that regulate core processes of neocortical circuit formation. This work further contributes to our understanding of the complex etiology of neurodevelopmental pathologies.

## RESULTS

### Satb2 controls neuronal migration and corpus callosum development cell-intrinsically

One of the most consistent features of Satb2 Associated Syndrome (SAS), and the Satb2 knockout mouse is agenesis of the corpus callosum^8, 10^. We previously found that layer II-III and layer V neurons use distinct molecular programs downstream of Satb2 to project axons to form this trans-hemispheric axonal tract ^13^. In order to dissect how Satb2 is required cell-autonomously for the axon development of late-born projection neurons, we used a targeted *Satb2* mouse strain where exon 2 of Satb2 is “floxed” (*Satb2^fl/fl^*) and is then deleted by cre-mediated recombination.

Satb2 distinctly orchestrates both cell extrinsic and cell intrinsic transcriptional programs important for this developmental process. When Satb2 is deleted from the developing dorsal neocortex using *Nex^Cre^*, Satb2-deficient neurons are sensitive to *cell extrinsic* effects and project their axons via the internal capsule, similar to what was observed in the Satb2 constitutive mutant ^13^ (Figure S1B).

However, when Satb2 is deleted in a mosaic fashion ie, *cell-intrinsically* in a wild-type cortex, neuronal migration is perturbed, and axons do not project at all (Figure S1C-E). Live imaging of these organotypic slices revealed distinct differences in the behavior of wild type and *Satb2-*deficient neurons in the intermediate zone (IZ, Figure S2). While wild-type neurons started migrating radially after acquiring a single leading process *Satb2*-deficient neurons failed to leave the IZ and often formed bifurcated leading processes.

Given *cell-intrinsic* neuronal migration and axon extension can be fully restored when re-introducing Satb2, while cell-extrinsic effects cannot (Figure S1), we sought to understand which Satb2 targets contribute to cell-intrinsic development of the corpus callosum.

### Sema7A acts downstream of Satb2 to control radial migration and axon elongation

In order to identify Satb2 downstream targets that control radial migration and axonal growth, we performed an *in situ* hybridization (ISH) screen with molecules known to be involved in axon formation and guidance (reviewed in ^29–31^). In this screen we used the *Nex^Cre^* mouse strain in order to delete Satb2 only in pyramidal neurons of the neocortex ^32^. We focused on genes expressed mainly in the cortical plate whose expression is altered in *Satb2^fl/fl^Nex^Cre^* brains at E18 as compared to wild type brains (Figure S3). We hypothesized that the expression of a certain ligand-receptor pair would be changed in Satb2 mutants since our previous experiments suggested both cell-intrinsic and cell-extrinsic (Figure S1) roles of Satb2 in neuronal migration and axon outgrowth. We reasoned that, the molecule acting as ‘receptor’ should be expressed in UL neurons where Satb2 has a cell autonomous role while its potential putative ‘ligand’ could be expressed in both deep layers and upper layers. We found one receptor-ligand pair that satisfied these criteria, Semaphorin7A/Integrinβ1 (Sema7A/Itgβ1). Expression patterns of both Sema7A and Itgβ1 were changed in the Satb2 mutant cortex. While Sema7A expression was reduced in UL neurons, Itgβ1 lost its lateral to medial gradient of expression within deep layers of the neocortex (Figure 1A). We also re-analyzed published RNAseq data from a different Satb2 mutant ^33^ and found that expression of Sema7A remains reduced at P0 (Figure 1B). In embryonic cortex, the Sema7A Transcription Start Site (TSS) and intron 1 are well marked by histone modifications associated with transcriptional activity such as H3K27ac, H3K9ac, and H3K4me1/3 (Figure 1C). ChIPseq peaks ^34^ for transcription factors NeuroD2, Tbr1 and Fezf2 have also been found near the TSS of Sema7A (Figure 1C).

**Figure 1.**
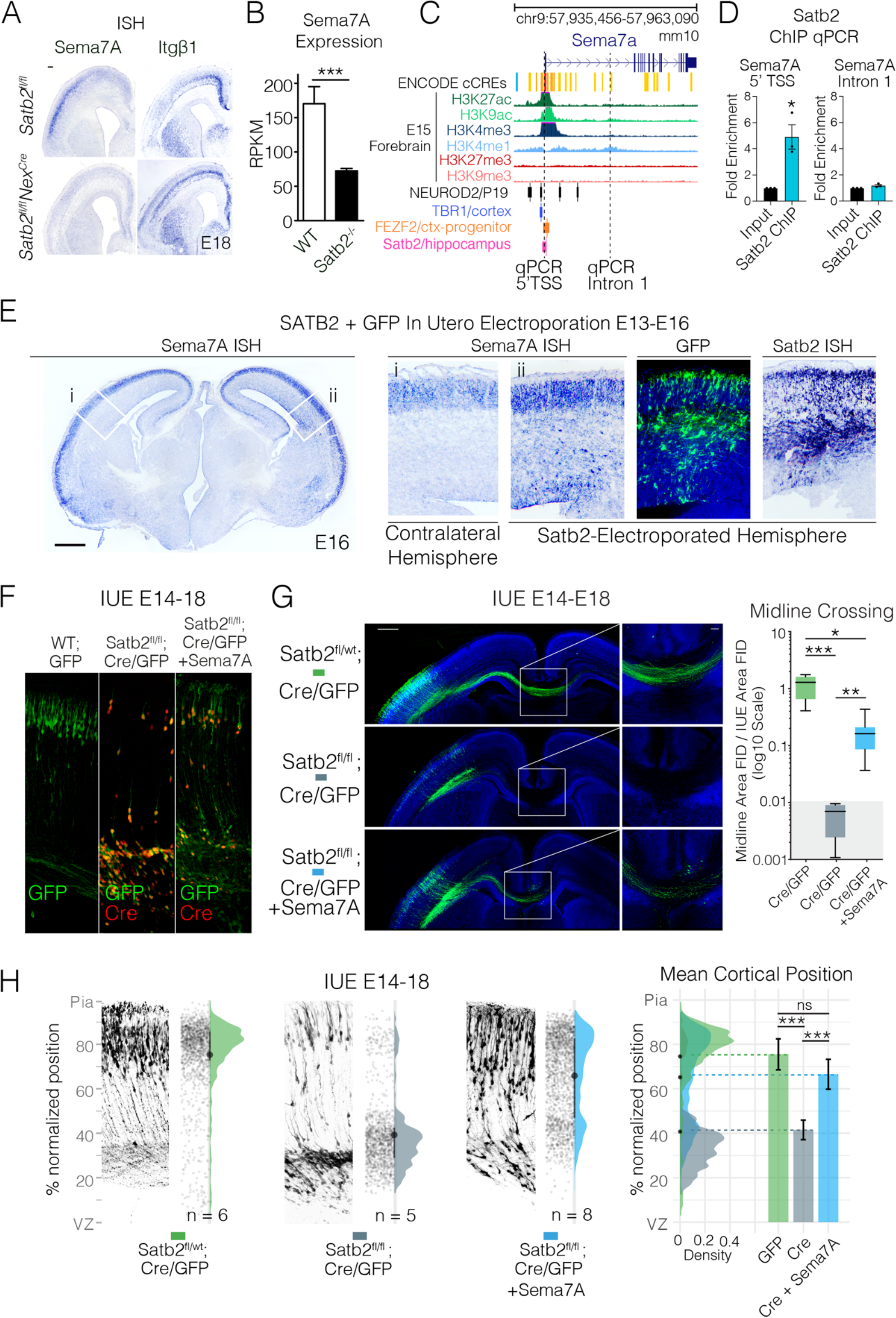
(previous page) Semaphorin7A is downstream of Satb2 and can restore migration and axon projection in Satb2 deficient neurons. (A) In situ hybridization (ISH) of *Sema7A* and Itgβ1 expression in the Satb2 cortex, scale bar is 100μm. (B) Sema7A expression at P0 quantified by RNAseq. (C) UCSC genome browser view of Sema7A gene locus with annotation tracks: ENCODE cis-regulatory elements (cCREs), Histone ChIPseq from embryonic forebrain and ReMap ChIPseq tracks for NeuroD2, Tbr1 and Fezf2, and Satb2-V5 Hippocampal ChIPseq ^36^ (D) Satb2 ChIP-qPCR of 5’TSS and Intron 1 Sema7A regions. (E) Ectopic *Sema7A* expression is observed after *Satb2* overexpression in *wild type* cortices. E16 wild-type brains electroporated at E13 with full-length *Satb2* full-length cDNA and GFP show higher expression of *Sema7A* RNA at the electroporation site (Scale bar 500 µm). (F) Overexpression of Sema7A into Satb2-deficient neurons restore migration cell-intrinsically. Immunostaining with GFP and Cre in *wild type* and *Satb2^fl/fl^* cortices after IUE with GFP (left), Cre/GFP (middle) and Cre/GFP +Sema7A (right). (G) Re-expression of *Sema7A* in *Satb2* deficient cells restores migration and partially rescues midline projections cell- intrinsically. *In utero* electroporations (IUE) into E14 embryos collected at E18 are shown with midline magnifications. In all conditions pNeuroD1- Cre + pCAG-FSF-GFP is abbreviated to Cre/GFP. Midline crossing was quantified by normalizing GFP Fluorescent Integrated Density (FID) at the midline to the Fluorescent Integrated Density of the electroporated area. Box and whisker plots represent Min-Max with lower and upper quartiles and means (right). [GFP] n = 5, [Cre] n = 4, [Cre + Sema7A] n = 8. Grey-ed out area at base of plot signifies measurements in this area are likely at the level of noise due to the complete absence of GFP fibers in [Cre] condition. Scale bar in the panoramic picture is 500μm, while scale bar in midline magnification is 100μm. (H) *Satb2* negative cells migrate into the cortical plate after *Sema7A* re-expression. Adjacent to example electroporation images are raw data points corresponding to cell positions, a half-violin showing the total cell distribution across all brains in that condition and mean (point) and interquartile range (line) between the raw data points and the half-violin. Cell distributions for the different conditions are shown overlaid on the right along with mean±SD cortical position. Statistics: qPCR in (D) used an unpaired t-test, midline quantifications (G) passed Shapiro-Wilk Tests for lognormality and log-transformed values were tested using one way ANOVA with Tukey’s multiple comparison. Migration profiles (H) passed Shapiro-Wilk test for normality and were tested using one way ANOVA with Bonferroni, ‘n’ displayed on figure refers to one electroporated cortex.

We performed Chromatin Immunoprecipitation using self-made Satb2 antibody ^35^ followed by quantitative real-time PCR (ChIP-qPCR) on E18 Cortical Lysates. We found Satb2 enriched in the 5’ TSS region and not in the H3K4me1-marked intron 1 of Sema7A (Figure 1C,D). This result was simultaneously confirmed in a Satb2-V5 ChIPseq ^36^ in adult hippocampus, where a Satb2 peak was observed at the TSS of Sema7A (Figure 1C, pink row). To test whether Satb2 expression is sufficient to induce ectopic *Sema7A* transcription, we expressed a *Satb2 cDNA* construct in the cortical VZ/SVZ at E13 by IUE (Figure 1E). This resulted in ectopic transcription of Sema7A mRNA at E16 in the VZ/SVZ as well as IZ regions of the neocortex, where it is not normally expressed (Figure 1E).

We then asked whether restoration of *Sema7A* expression in *Satb2*- deficient neurons could rescue defects of axonal specification and/or neuronal migration. Immunostaining showed that all GFP expressing cells also expressed Cre but lacked Satb2 expression (Figure 1F). Re-expression of *Sema7A* largely enabled midline crossing of Satb2-deficient axons *in vivo* (Figure 1G). It also restored the laminar position of Satb2- deficient neurons (Figure 1H). Our results show that Sema7A is a direct *Satb2* target that promotes radial neuronal migration and axon outgrowth during neocortical development.

### Sema7A is required for polarity acquisition and axon outgrowth

There are three crucial steps in the development of callosal projections: the specification of the axon and the leading process, axonal extension, and midline crossing. It is widely accepted that in the neocortex, onset of axon extension coincides with initiation of neuronal migration when a multipolar cell becomes polarized ^2, 37^. To better understand the phenotype of Satb2-deficient neurons (Figure S1), we cultured dissociated Satb2-deficient (Satb2-/-) and wildtype (WT) neurons *in vitro* and characterized their morphology (Figure 2). Future axons were identified as the longest neurite at DIV2 and by counterstaining for the axonal marker TAU at DIV4.

**Figure 2.**
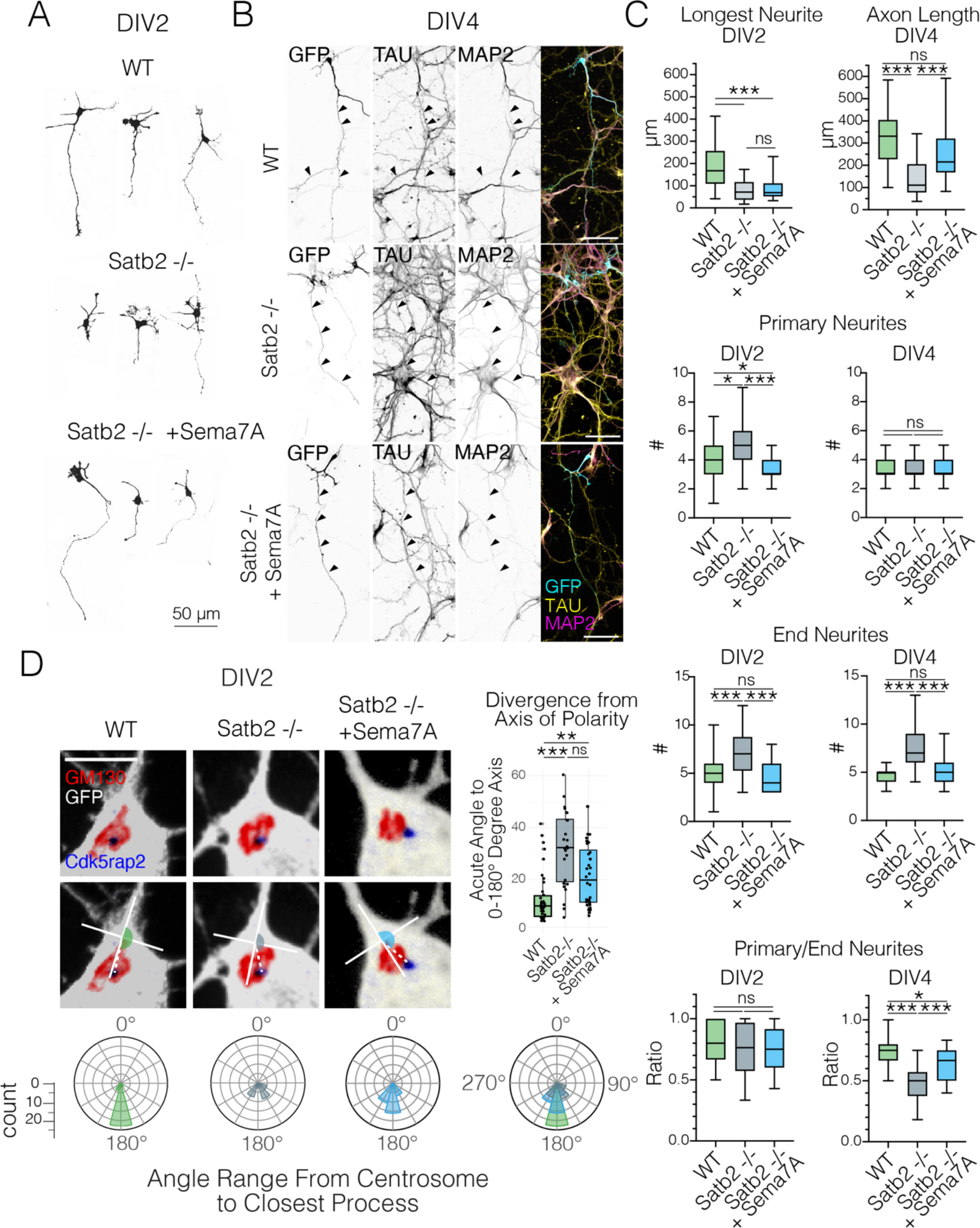
(Previous Page). Sema7A restores the polarization and axon outgrowth in *Satb2-*deficient neurons also *in vitro*. (A) Examples of WT, Satb2 deficient, and Sema7A rescue conditions in E14 primary cortical neurons after 2 days in vitro (DIV2). (B) After 4 days in vitro (DIV4), axons can be identified in all conditions by their enrichment of TAU-1 and depletion of MAP2. Scale bar=50μm. (C) Quantification of neuronal morphology at DIV2 (left column) and DIV 4 (right column). All box & whisker plots in C plot min, max, interquartile range and median. Data were non-normally distributed (D’Agostino & Pearson, Shapiro-Wilk tests). For all tests, Kruskal Wallis test followed by Dunn’s Multiple comparison was used, where adjusted p values < 0.001 = ***, < 0.01 = **, < 0.05 = *. DIV2 WT n= 48, Satb2 -/- n = 54, Satb2-/- + Sema7A n = 37; DIV4 WT n = 20, Satb2 -/- n = 37, Satb2-/- + Sema7A n = 39 cells. (D) Loss of Satb2 is associated with a random distribution of the centrosome which can be restored by Sema7A. (Top row): Representative images of DIV2 primary cortical neurons stained with the golgi marker Gm130 and centrosomal marker Cdk5rap2 (scale bar 50μm). Radar plots depict the circularized histograms of angle counts, where 360 degrees are binned in 30-degree increments. Measured blind using a standardized 100 x 100 pixel cross, which was placed with the parallel axis centered in the process and the perpendicular access touching the Golgi. Data were non-normally distributed (D’Agostino & Pearson, Shapiro-Wilk tests). For all tests, Kruskal Wallis test followed by Dunn’s Multiple comparison was used, where adjusted p values < 0.001 = ***, < 0.01 = **, < 0.05 = *. WT n=35, Satb2-/- n=25, Satb2-/- + Sema7A n = 35.

We observed that while Satb2-/- neurons do project axons in vitro, these axons are markedly shorter than WT cells at DIV2 and DIV4 (Figure 2B,C). Furthermore, Satb2-/- cells possess more primary neurites than WT cells at DIV2, which appears to result in increased branching (End neurites) by DIV4 (Figure 2A,C). While Satb2-/- neurons do project axons *in vitro*, these axons are markedly shorter than WT and these neurons do not achieve complete polarization (Figure 2B,C). Sema7A re-expression restored the number of primary and end neurites to WT levels (Figure 2A,C). By DIV4, most Sema7A-rescued Satb2-/- neurons also showed recovered TAU+/MAP2- axon lengths.

Previous studies have reported that a young neuron becomes polarized when the microtubule organizing center, the centrosome, positions itself together with the Golgi apparatus in front of the neurite that will become the axon, and prior to migration moves to specify the leading process ^38–40^. To examine the possible role of Satb2 in mediating polarization, we measured the positions of the centrosome and Golgi organelles with respect to the closest process. At DIV2 in wild type neurons, the centrosome lies at the base of and in line with the longest neurite along the 0-180° ‘polarity’ axis (Figure 2D, left). Loss of Satb2 was associated with a disturbance in the position of the centrosome was disturbed and this structure was mostly found between neurites, resulting in angles that are further away from the polarity axis (Figure 2D, middle). We then analyzed the role of Sema7A in regulating neuronal morphology and polarity downstream of Satb2 by overexpression of Sema7A in these cells. We found that re-expression of *Sema7A* in *Satb2 -/-* neurons shifted the centrosome angle back towards the polarity axis (Figure 2D, right). Collectively, our results show that Sema7A controls neuronal polarity and outgrowth as well as initiation of both migration *downstream of Satb2*.

### Sema7A Membrane localization and Dimerization are required to mediate cell-intrinsic effects

Since Sema7A lacks a cytoplasmic domain and is reported to act as a ligand *in trans* ^41^, it was difficult to envisage how it mediated a cell-intrinsic rescue in Satb2 deficient neurons. One possibility was that Sema7A was being secreted to act in an autocrine fashion on the cell expressing it. We therefore repeated genetic rescue experiments with a mutant form of Sema7A, where we removed the GPI membrane anchor (ΔΜΕΜ). Expression of the secreted version of Sema7A (ΔΜΕΜ) was neither capable of restoring migration, nor axon elongation in Satb2 mutants (Figure 3B, C). This indicates that Sema7A requires membrane attachment in order to function cell intrinsically downstream of Satb2. Another possibility is that Sema7A acts by complexing in *cis* with other receptors, most likely transmembrane proteins capable of reverse signaling. The seven-bladed beat-propeller SEMA domain has been shown to mediate dimerization of Semaphorins, a characteristic essential for their function ^42, 43^, as well as interaction with other receptors (eg Nrp1, ^44^). Deletion of this important domain also interferes with Sema7A function downstream of Satb2 (Figure 3B, C) indicating that dimerization is essential for Sema7A function. We also generated a version of Sema7A where we mutated the conserved Arginine Glycine Aspartate (RGD) motif required for Integrin binding ^45^, (KCE mutant, Figure 3A). Mutation of this site interestingly disrupted axon outgrowth but not migration. We obtained a similar result upon deletion of the structural IG or the PSI domain that in other SEMA members is required for the correct positioning of the ligand-binding site^46^, suggesting differences in the signaling roles of Sema7A during these two processes. Together these data show that Sema7A is acting as a membrane-associated receptor most likely in complex with other receptors.

**Figure 3.**
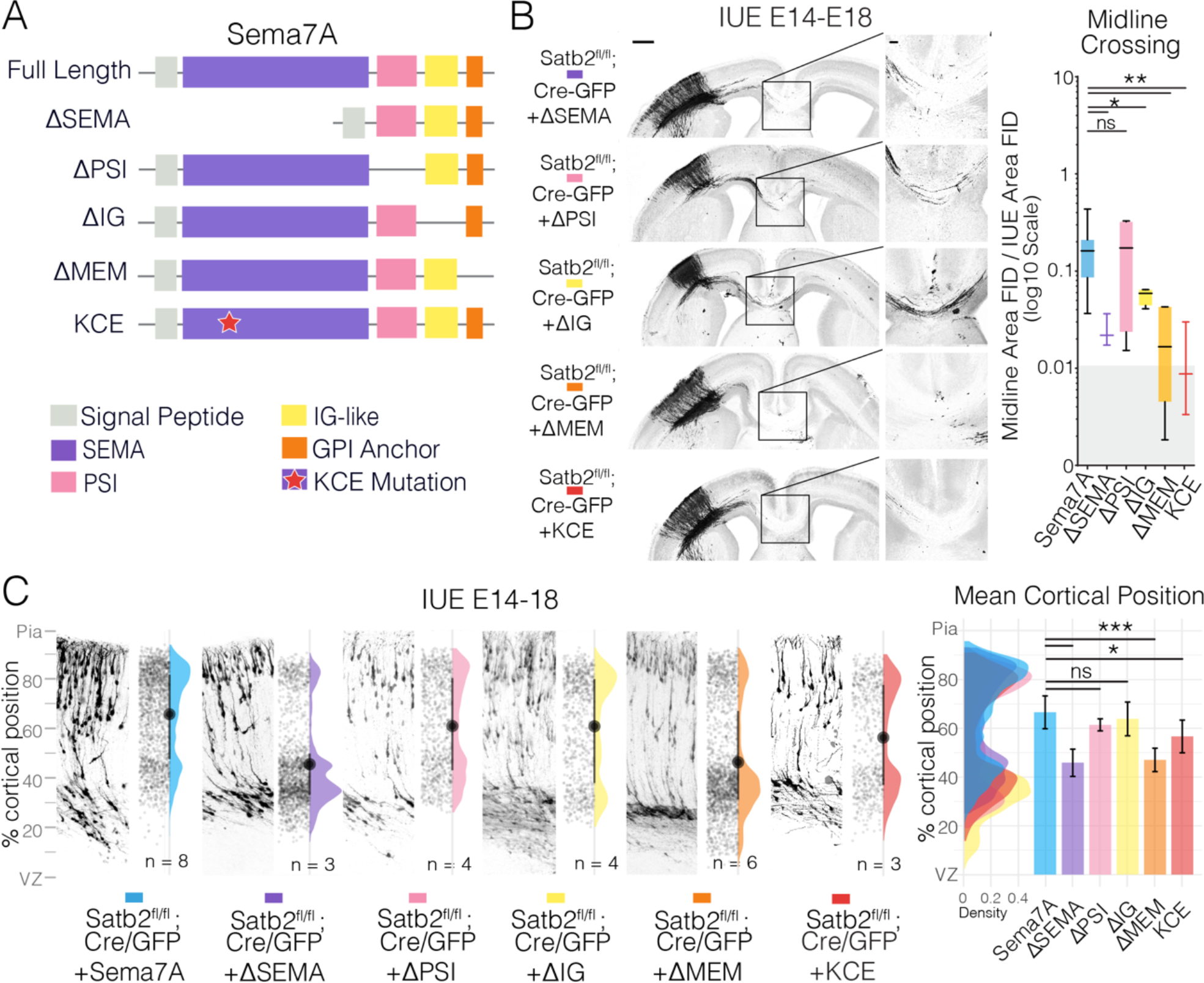
**Semaphorin 7A Domains associated with membrane localization and dimerization are required for migration and axon outgrowth**. (A) Schematic of Sema7A domains and deletion mutants. The scheme shows all currently annotated domains of Sema7A and depicts the deletion constructs generated. Sema domain is depicted in purple, the Plexin-Semaphorin-Integrin (PSI) domain in pink, the immunoglobulin domain (IG) in yellow and the Glycosyl-Phosphatidyl-Inositol (GPI) membrane anchor is depicted in orange. The Arginine-Glycine-Aspartic acid (RGD) reported to be important for Integrin binding was mutated to Lysine-Cysteine-Glutamic Acid (KCE) and is depicted as a red star. (B) Panoramas of *Satb2^fl/fl^* IUE brains with the indicated Sema7A deletion constructs together with pCAG-FSF-GFP and pNeuroD1-Cre and midline magnifications and quantifications. Midline crossing is quantified as described previously. Box and whisker plots represent min-max with lower and upper quartiles and means. [Cre + Sema7A] (shown in Figure 2) n = 8, [Cre + Sema7A-ΔSEMA] n = 3, [Cre + Sema7A-ΔPSI] n = 4, [Cre + Sema7A-ΔIG] n = 4, [Cre + Sema7A-ΔMEM] n = 6, [Cre + Sema7A-KCE] n = 3. Grey-ed out area at base of plot signifies measurements in this area are likely at the level of noise due to the complete absence of GFP fibers in [Cre] only condition (Figure 2). Scale bar in panoramic picture is 500 μm, while scale bar in midline magnification is 100 μm. (C) (Left): Sema7A deletion mutant migration profiles from E14-18 IUEs. Each subplot is comprised of a representative electroporation image on the left along with raw data points, and a half violin plot marked with the mean (dot) and interquartile range (line). (Right): Cell distribution plots distributions are overlaid on the right along with mean±SD cortical position. *Statistics:* data in B were lognormally distributed and were log-transformed prior to running a one way ANOVA with Bonferroni multiple comparison. In C, Kruskal-Wallis with Dunn’s multiple comparison.

### Sema7A acts as a co-receptor of Sema4D

We asked whether Sema7A could heterodimerize with other semaphorin family members that could transmit signals into the expressing cell in a cell autonomous manner. Members of Semaphorin Class 4 and Class 6 had recently been been shown to be involved in reverse signaling during cell polarization and migration ^24–27^. We focused on semaphorins that contain a cytoplasmic domain and could thereby transmit signals into the expressing cell in a cell autonomous fashion. We also reasoned that the expression should overlap with that of Sema7A, and should remain in Satb2 deficient cortex (Figure S3). We first performed Co-Immunoprecipitation (Co-IP) in HEK293T cells using a C-terminal GFP tagged Sema7A and N-terminal Myc-tagged versions of Sema5A, Sema6A, and Sema4D – three putative receptors that meet above mentioned criteria (Figure 4A). We detected a strong binding affinity between Sema7A and Sema4D and a weak interaction with Sema6A (Figure 4A). Notably, Sema4D is also expressed in the developing cortical plate alongside Sema7A (Figure 4B). While Sema7A is reduced in the Satb2 mutant, Sema4D is modestly upregulated (Figure S2).

**Figure 4.**
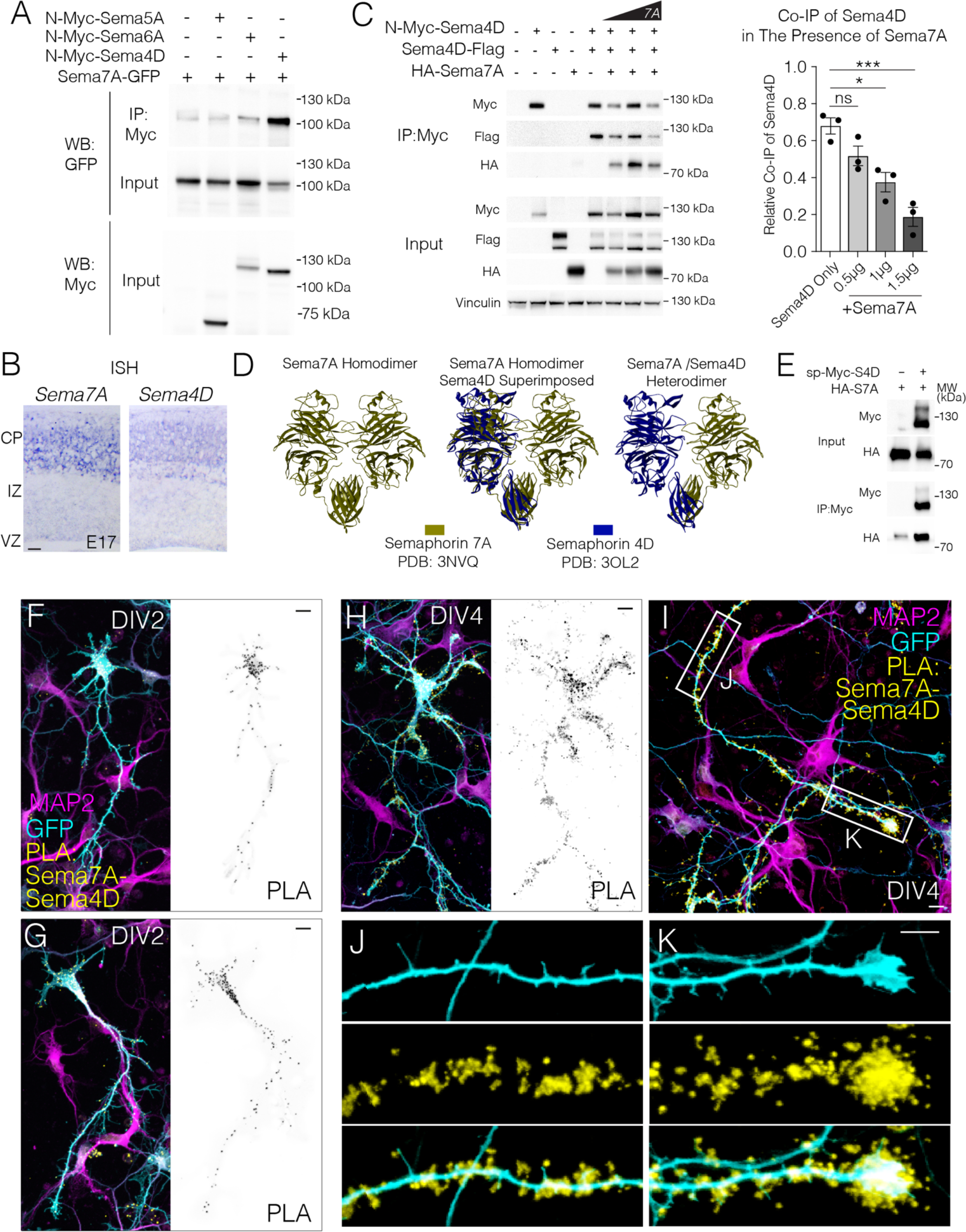
(previous page). Sema4D binds with high affinity to Sema7A. (A) Sema7A binds to Sema4D and Sema6A. HEK293T cells were transfected with the GFP tagged Sema7A (this paper) and the indicated Myc-tagged Sema family members: *N-*Myc-Sema4D (addgene # 51599, ∼120kDa), N-myc-Sema5A (∼65kDa), and N-myc-Sema6A (∼120/130kDa). (B) Competition assay reveals Sema4D homodimerization decreases with increasing concentrations of Sema7A. (C) *In Situ Hybridization* (ISH) at E18 in wild type mouse cortex shows the expression of both Semaphorins in the cortical plate. Scale bar= 100μm. (D) Co-immunoprecipitation of HA-Sema7A using sp-myc-Sema4D. (E) Heterodimer alignment of Sema4D and Sema7A using published crystal structures. (F-K) Localization and distribution of Sema4D-Sema7A complexes in primary cortical neurons. Primary E14 neurons transfected with sp-Myc-Sema4D, HA-Sema7A, and GFP were fixed at DIV2 and DIV4 and subcellular localization of semaphorin complexes determined by Proximity Ligation Assay (PLA). GFP signal (cell fill) can be seen in cyan, Sema4D-Sema7A PLA signal in yellow, and MAP2 in magenta. Scale bars in F-K are 10μm.

Sema4D is known to form homodimers^47^. To test whether Sema7A competes with Sema4D for dimerization, we additionally cloned a version of Sema7A that contains an HA tag at an exposed loop of the Sema domain at codon 352, (HA-Sema7A) and a Sema4D with a C-terminal (cytoplasmic) flag tag. Increasing amounts of Sema7A resulted in decreased amounts of Sema4D-Flag co-immunoprecipitated with N-Myc- Sema4D, indicating that Sema7A competes with Sema4D monomers for dimerization (Figure 4C). We observed 150kDa and 120kDa forms of Sema4D similar to previous reports^48^. 120kDa and 150kDa forms of Sema4D are both ‘full length’ given the detection of the C terminal tag, however, in the presence of N-Myc-Sema4D, Sema4D-flag is observed more prominently as the 120kDa form, which is also the preferred form pulled down by co-immunoprecipitation in flag lysis buffer (Figure 4C). This suggests the existence of differential post-translational modification of the homodimerized form of Sema4D as compared to Sema4D present in Sema4D-Sema7A heterodimers.

Given N-Myc-Sema4D (addgene #51599) uses a chimeric signal peptide and produces very little of the 150kDa form of Sema4D ^49^, we cloned Sema4D to contain a myc tag after its endogenous signal peptide based on structural data (termed sp-Myc-Sema4D). This construct successfully yields both ∼120kDa and 150kDa forms of Sema4D (Figure 4D).

We used the published crystal structures of Sema4D ^50^ and Sema7A ^41^ to model the Sema7A:4D heterodimer. We found that superimposition of one Sema4D molecule over the one Sema7A molecule from the published Sema7A homodimer, resulted in only a minor deviation (RMSD:0.64 Å) in the C^α^ atoms of the proteins (Figure 4E), further supporting the existence of a physiological competitive interaction between these two proteins.

To visualize the interaction of Sema7A with Sema4D *in vivo*, we carried out a Proximity Ligation Assay (PLA) in wild type cortical neurons transfected with sp-Myc-Sema4D and HA-Sema7A (Figure 4F-K). Indeed, we observe a direct interaction of HA-Sema7A and Sema4D in neurons. At DIV2, the Sema4D-Sema7A heterodimer can be seen in the soma and axon hillock (Figure 4F,G). In more mature neurons at DIV4, PLA signal is further enriched in the growing axon and is observed at the tips of filopodia, branch points and with an enrichment at axonal growth cones (Figure 4H-K).

### Sema4D is required for cell-intrinsic Sema7A-mediated neuronal migration and axonal growth

To address whether Sema7A and Sema4D function together in promoting neuronal migration and axonal growth cell-intrinsically, we conducted loss-of-function experiments in vivo. We first addressed whether the role of Sema7A downstream of Satb2 in promoting migration and axon outgrowth was dependent on the function of Sema4D. This was carried out by downregulating Sema4D expression using a specific shRNA, while restoring Sema7A expression in Satb2-deficient neurons in vivo.

As previously observed, re-expression of Sema7A improved the laminar position (Figure 5A) and callosal projections of Satb2-deficient neurons (blue condition, Figure 5B,C). Downregulation of Sema4D, however, prevented Sema7A-mediated rescue of both migration (Figure 5A) and axonal outgrowth in the *Satb2*-deficient neurons (Figure 5B-C, red condition).

**Figure 5.**
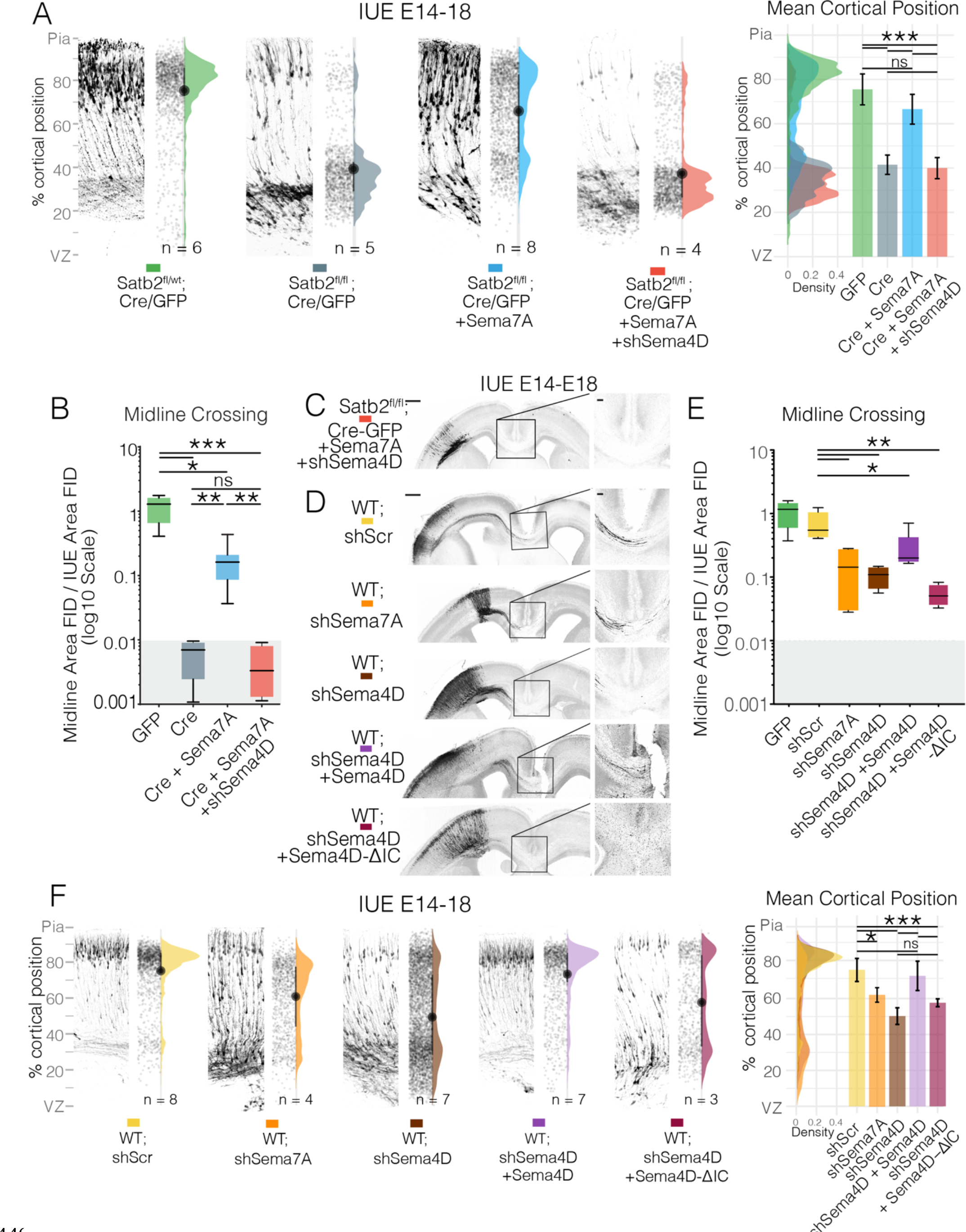
(previous page). The Cytoplasmic Domain of Sema4D is required for cell intrinsic migration and axon outgrowth. (A) Simultaneous downregulation of Semaphorin 4D by shRNA reverses Semaphorin 7A rescue of neuronal migration in Satb2-deficient neurons. IUE into E14 embryos collected at E18. Cell distributions are overlaid on the right along with mean±SD cortical position. (B) Quantification of midline crossing after simultaneous downregulation of Semaphorin 4D in Sema7A-rescued Satb2-deficient neurons with example depicted in (C). Panoramas of Sema4D and Sema7A shRNA electroporations with midline magnifications. Midline crossing of shRNA electroporation (D) is quantified as described previously. Scale bar in panoramas is 500 μm, while scale bar in midline magnification is 100 μm. Box and whisker plots (B, C) represent Min-Max with lower and upper quartiles and means. Midline [GFP] n = 5, [Cre] n = 4, [Cre + Sema7A] n = 8, [Cre + Sema7A + shSema4D] n = 4 [shScr] n = 5, [shSema7A] n = 4, [shSema4D] n = 4, [shSema4D + Sema4D] n = 7, [shSema4D + Sema4D-ΔIC] n = 4. (F) Migration profiles of neurons after downregulation of Sema4D. Control scrambled shRNA (shScr), shRNA against Sema4D, shRNA against Sema4D + Sema4D cDNA, or shRNA against Sema4D plus a version of Sema4D cDNA lacking the intracellular domain -IC (Sema4D-ΔIC). Representative electroporations are shown adjacent to raw data points and half violin plots of the total distribution with mean±SD cortical position on the right. *Statistics*: (A,B,E) one way ANOVA with Bonferroni, values in (B & E) were lognormally distributed and log-transformed prior to test. (F) Kruskal-Wallis test with Dunn’s Multiple comparison.

We also tested whether downregulation of Sema4D or Sema7A by shRNA would phenocopy cell autonomous deletion of Satb2. In a similar design, we introduced a construct expressing *shRNA* against Sema7A into E14 cortex and analyzed the brains at E18. Indeed, Sema7A knockdown resulted in reduced projection of axons into the contralateral hemisphere (Figure 5D,E) and a migration deficit of upper layer neurons (Figure 5F). Similar experiments using shRNA against Sema4D, produced even stronger phenotypes, with strong disruption of both migration and axon growth (Figure 5D-F).

Our hypothesis of Sema7A was based on its interaction in cis with a transmembrane receptor that could thereby mediate reverse signaling. We therefore addressed if the intracellular domain (ICD) of Sema4D is required for it function in neuronal migration and axonal growth. To test this, we downregulated Sema4D using a specific shRNA, and at the same time overexpressed either full length Sema4D, or Sema4D lacking the ICD (Sema4D-ΔIC). Only the construct encoding the full cDNA could rescue defects of both migration and axon outgrowth caused by Sema4D loss-of-function, while Sema4D-ΔIC was unable to repair these defects (Figure 5D-F, light and dark purple conditions). Together, these data show that the intracellular domain of Sema4D is essential for the cell-autonomous effects of Sema7A on the radial migration and axonal outgrowth of upper layer neurons in the developing neocortex.

### Human Sema4D-497P mutation inhibits trafficking to the membrane and growth cone

Over the course of this study, we identified a patient presenting with generalized tonic-clonic seizures that has a de novo mutation in Sema4D (see case report in supplemental note 1). Exome sequencing revealed a non-synonymous mutation of adenosine to cytosine resulting in a glutamine (Q) to proline (P) substitution at codon 497 (Figure 6A). Given that the Q497 residue is involved in the formation of a beta sheet on a propeller of the sema domain (Figure 6B), the sudden inclusion of a proline ring was expected to alter secondary structure. To predict how this mutation may affect protein folding, we used the most recent release of alphafold2 ^51^ to compare predicted structure with the known crystal structure of hSema4D (PDB: 1OLZ). Interestingly, the predicted structure of wildtype hSema4D (Figure 7B, middle panel) and hSema4D-497P are extremely close to the 1OLZ crystal structure (Figure 6B, upper panel). In both WT and 497P predictions, upper segments of the sema domain form low confidence loops, due to a limitation of alphafold2 (Figure S4). However, the beta sheet containing codon 497 is consistent between 1OLZ and the prediction for hSema4D, so the comparison between 497 mutation structure to wildtype is useful at this level. While the beta sheet containing P497 is predicted to form correctly, the beta sheet below this is absent in the predicted model of hSema4D-Q497P. This lower beta sheet normally resides in close proximity to glycosylation site N77 (Figure 6B, lower panel zoom). It is also possible that steric hindrance from the Q497P mutation may affect normal glycosylation at N49 and N419 or ubiquitination at K505 or K81 (Figure 6B, lower panel zoom).

**Figure 6.**
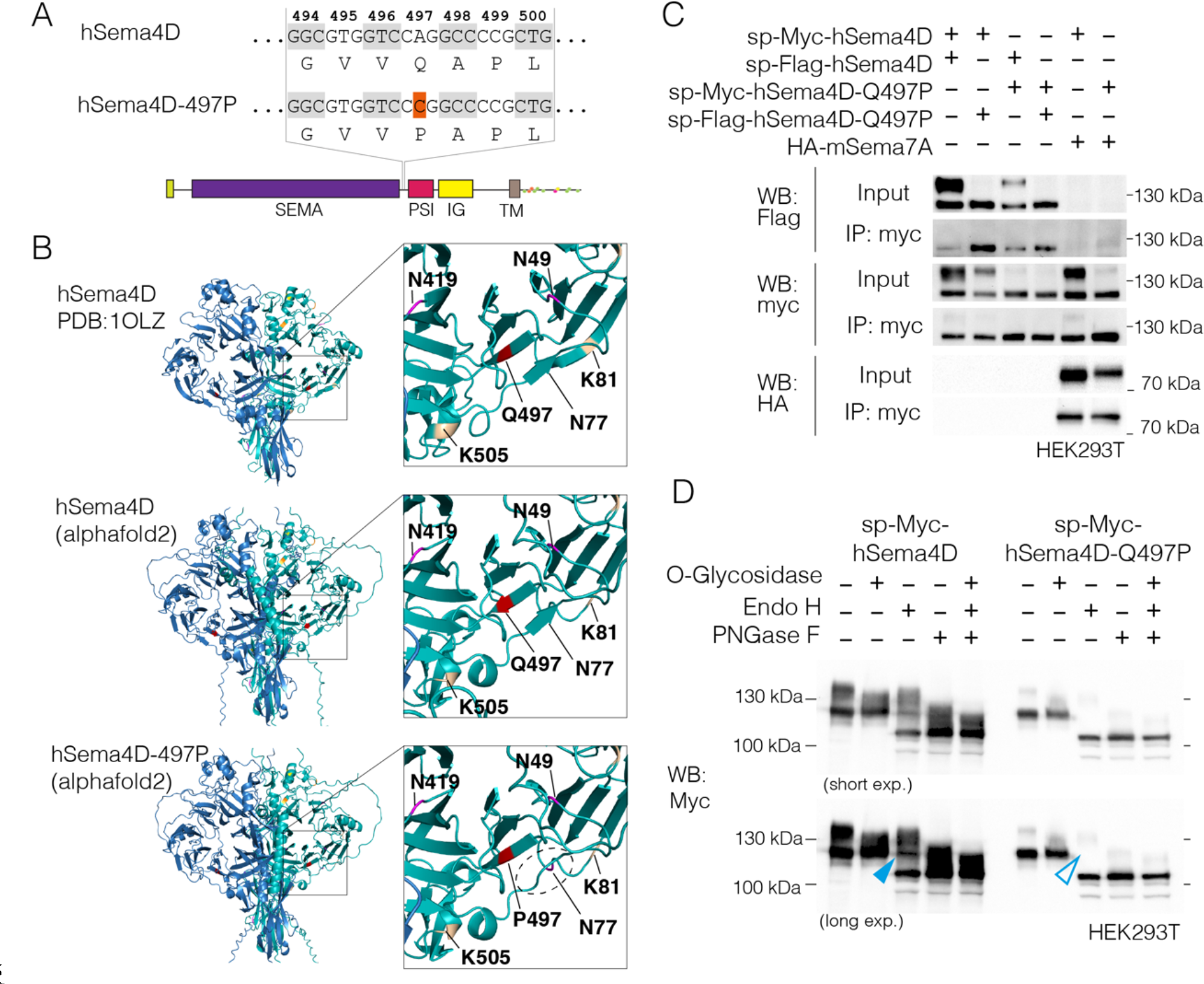
(previous page). Human Sema4D-497P Mutation retains homo and heterodimerization ability but is improperly processed. (A) Schematic of adenosine to cytosine base mutation giving rise to a glutamine ‘Q’ to proline ‘P’ substitution at codon 497 between SEMA and PSI domains. (B) Ribbon diagrams and zooms of hSema4D solved crystal structure (1OLZ, top) with alphafold2 predictions of wildtype hSema4D (middle) and hSema4D with 497P mutation (bottom). Codon 497 is highlighted red, glycosylation residues magenta, phosphorylation residues yellow, and ubiquitination residues ochre. A beta sheet parallel to codon 497 (dotted ellipse) is not predicted to form following 497P mutation. (C) Co-immunoprecipitation of myc-tagged human Sema4D (sp-myc-hSema4D) and the 497P variant (sp-myc-hSema4D- Q497P) confirms that the mutant form can still form homodimers with sp-flag- hSema4D and heterodimers with HA-mSema7A. While hSema4D produces bands at ∼150kDa and ∼120kDa, the hSema4D-Q497P variant predominantly generates the 120kDa band. (D) De-glycosylation assay of sp-myc-hSema4D and sp-myc- hSema4D-Q497P using O-Glycosidase, Endo H and PNGase F. The same blot is shown at short and long exposures.

**Figure 7.**
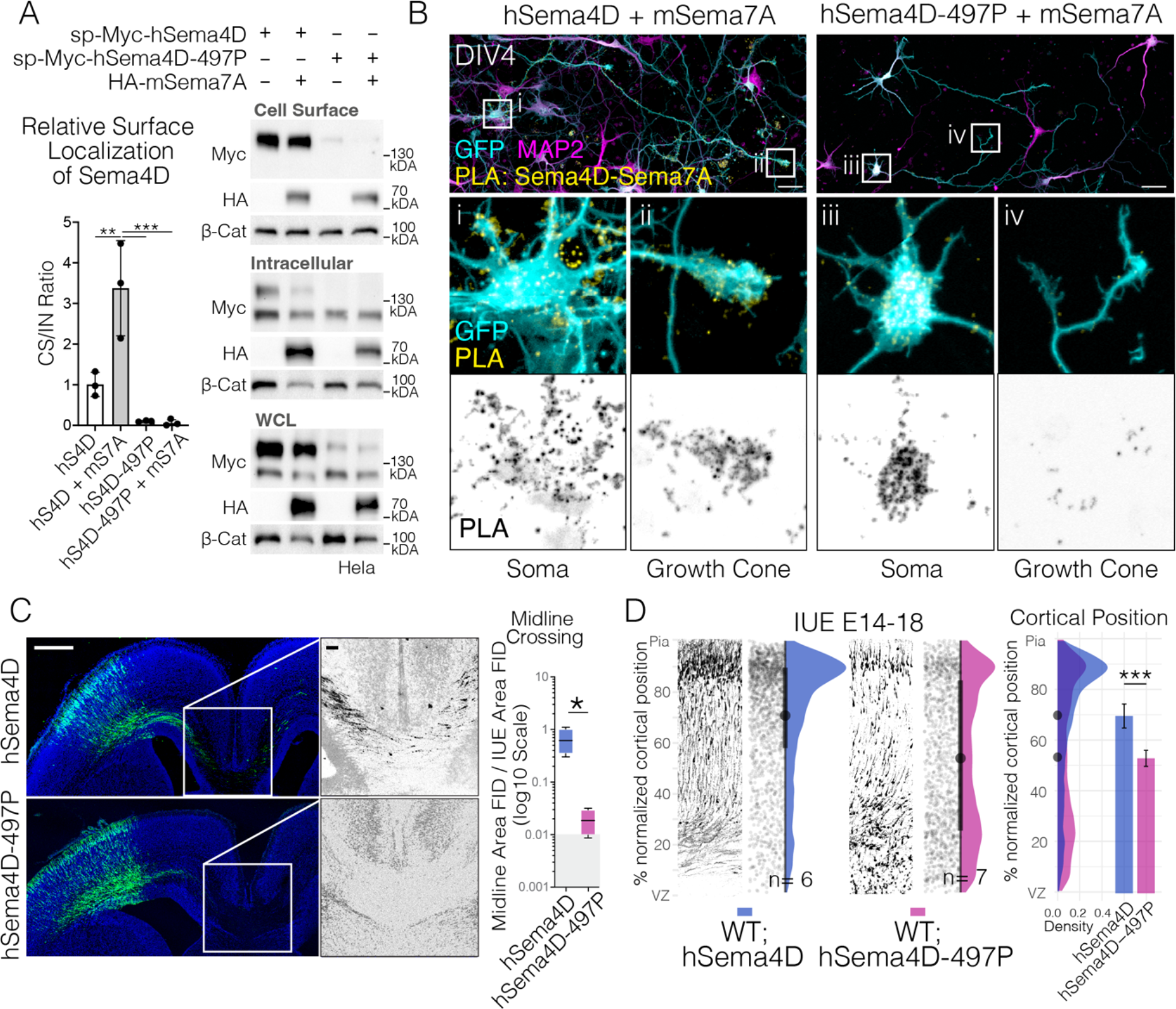
497P Mutation abolishes localization of hSema4D and Sema7A to the cell surface and growth cones and reduces neuronal migration and axon projection in vivo. (A) Surface biotinylation followed by avidin pull down of sp-myc- hSema4D or sp-myc-hSema4D-497P in the presence or absence of HA-mSema7A. Cell Surface to Intracellular (CS/IN) ratio of Sema4D was calculated by first normalizing each fraction to all Sema4D detected in whole cell lysate (WCL) where the complete calculation is CS/IN = (CS/WCL)/(IN/WCL). Plotted is mean+SD, using n = 3 separate experiments. (B) Proximity Ligation Assay (PLA) detecting interaction between mouse HA-Sema7A and human sp-myc-Sema4D or human sp-myc- Sema4D-497P. (C) Midline panoramas and quantifications of human Sema4D + GFP (hSema4D) and human Sema4D-497P + GFP in utero electroporations. Midline fluorescence was quantified as before, fluorescence intensities are log-normally distributed, so were log-transformed prior to running an unpaired t-test with Welch’s correction. hSema4D n = 4 sections over two biological replicates, hSema4D-497P n = 4 over 3 biological replicates. Scale bar in panoramas is 500 μm, while scale bar in midline magnification is 100 μm. (D) Migration profiles of cortical neurons following in utero electroporation with hSema4D or hSema4D-497P. hSema4D n = 6 sections over two biological replicates, hSema4D-497P n = 7 sections over 3 biological replicates.

We asked if Sema4D-497P can still form homo- and heterodimers after immunoprecipitation of proteins overexpressed in HEK293T cells. The Q497P mutation did not interfere with Sema4D homodimerization or heterodimerization with Sema7A (Figure 7C). However, we observed a shift of the Sema4D bands in an SDS-PAGE gel. While wildtype Sema4D normally migrates as two sizes (∼150kDa and ∼120kDa), the heavier form of the protein (∼150kDA) was largely absent in Sema4D-497P (Figure 6C).

In line with our structural predictions, we hypothesized that the two forms of Sema4D could correspond to different maturation states or glycosylation states of the protein. To test this hypothesis, we incubated hSema4D overexpressing lysates with de-glycosylation enzymes. De- glycosylation with O-Glycosidase resulted in the loss of the 150kDa migrating form of hSema4D and the presence of only a 120kDa form, indicating that this higher migrating form results from *O*-linked glycosylation of the protein. Treatment of hSema4D with PNGase F, which cleaves all glycosylation types, results in further downwards shift of the Sema4D band to 110kDa, indicating additional N-glycosylation is present on Sema4D. Endo H, which cannot cleave complex glycans, results in a mixed picture of 150, 120 and 110kDa migrating forms of hSema4D.

We then analyzed glycosylation on the mutated form of hSema4D. hSema4D-Q497P runs at 120kDa and lacks the 150kDa migrating form that was associated with O-linked glycosylation in wildtype hSema4D. Indeed hSema4D-Q497P was insensitive to O-Glycosidase indicating that unlike the wildtype form, hSema4D-Q497P lacks O-linked glycosylation. hSema4D-Q497P is still sensitive to PNGase F, showing that the protein is nevertheless modified by other glycosylation types. The uncomplete glycosylation of hSema4D-Q497P is also evident upon incubation with Endo H, which only yields a 110kDa form, compared to the intermediate 120kDa fragments observed when digesting WT hSema4D (Figure 6D, WT blue arrow vs Q497P empty blue arrow).

Notably however, treatment with PNGase F, which should cleave all glycosylation, still results in two prominent WT bands, reduced to ∼120kDa and ∼110kDa, suggesting that something additional to glycosylation contributes to the difference in protein size.

Given that presence of the Sema7A receptor at the plasma membrane is essential for its biological function, and glycosylation is known to be important in protein localization^52^, we used surface biotinylation followed by avidin pull down to address the role of Sema7A and of the Q497P mutation in regulating the subcellular localization of Semaphorin 4D (Figure 7A). Interestingly, only the 150KDa form of Sema4D became biotinylated, indicating that this O-linked glycosylated form is the membrane-localized form of the protein. We observed that co-expression of Sema7A increased the proportion of surface-localized Sema4D three-fold, while mutation of Sema4D at residue 497 abolished its localization to the cell surface.

To further characterize this effect, we nucleofected primary neurons with HA-mSema7A and sp-myc-hSema4D or sp-myc-hSema4D-Q497P and performed a proximity ligation assay to observe the location of the semaphorin complex in situ (Figure 7B). We observed that while hSema4D-mSema7A complexes are normally found on the outer side of the cell membrane and the growth cone (as per Figure 4), hSema4D- 497P-Sema7A complexes are predominantly found intracellularly the soma and absent from the growth cone (Figure 7Bi-iv), indicating loss of Sema4D-Sema7A complexes from the plasma membrane.

Finally, we analyzed the functional consequences of mutation of Sema4D during neocortical development. Overexpression of hSema4D- Q497P in the developing neocortex by *in utero* electroporation, disrupted radial migration and neurons collected in the lower portions of the neocortex (Figure 7C,D). Furthermore, overexpression of hSema4D- Q497P prevented axon projection across the corpus callosum (Figure 7C). Thus mutation of Sema4D at Q497 phenocopies downregulation of either Sema7A or Sema4D. Given that the mutation does not interfere with Sema4D homo- or heterodimerization (with Sema7A), together this data suggests that the dominant negative actions of this mutation results from defective O-linked glycosylation events and ensuring failure of Sema4D-Sema7A complex localization to the plasma membrane.

In this study we observed that a direct target of Satb2, Semaphorin7A, can mediate potent cell-autonomous signaling that facilitates neuronal migration and axon projection in the developing cortex (Figure 1). Membrane localization of Sema7A is essential for this function as is interaction of this GPI-linked protein with Sema4D, a transmembrane Semaphorin family member. Sema7A-Sema4D complexes can be found at neurites and are enriched in the growth cone (Figure 4). Membrane localization of the Sema7A-Sema4D complex is also dependent on correct O-linked glycosylation of Sema4D. We identified a human mutation in Sema4D, Q497P associated with tonic-clonic seizures, that interferes with the membrane localization of the Sema7A-Sema4D complex by disrupting most likely O-linked glycosylation events on Sema4D. This mutant acts in a dominant negative fashion to interfere with neuronal migration and axon projection during neocortical development.

## DISCUSSION

Here, we characterized a potent role of the Satb2-target Sema7A, in mediating cell-intrinsic migration and axon elongation in cortical projection neurons. Sema7A is expressed robustly in newly born neurons, and in addition to being a Satb2- downstream gene, Sema7A appears to be a target of several early neuronal transcription factors such as NeuroD2, Tbr1, and Fezf2 (Figure 1). Previous studies have identified Sema7A to be important for olfactory bulb, olfactory- and thalamo-cortical tracts, as well as for hippocampal and cortical axonal outgrowth and branching ^21, 45, 53^. However, in these studies, Sema7A was considered a classical ligand that initiates signaling cascade in a neighboring cell, after binding to its receptor Integrinβ1 or PlexinC1. The interpretation of these previous studies may need to be adjusted to account for Semaphorin- Semaphorin interactions and reverse signaling; of three Semaphorins tested, in addition to the strong interaction between Sema7A and Sema4D, we could detect weak binding between Sema7A and Sema6A. Further experiments are needed to characterize the full extent of Semaphorin-Semaphorin interactions and their capacity to carry out reverse signaling.

It is known that some Semaphorins can homodimerize ^19, 23^ and act as ligands which bind *in trans* to their respective receptors ^28, 53^. Here, we show the interaction of the two immune Semaphorins, Sema4D and Sema7A, that heterodimerize (Figure 4). By downregulating each of them independently, we observe disruption in the radial neuronal migration and less axons crossing the midline. Moreover, by simultaneously knocking down Sema4D in the Sema7A rescue condition, we show that migration and axon defects return (Figure 5). Thus, Sema4D is required for the Sema7A- mediated rescue in Satb2-negative cells. Additionally, our computational protein structure model suggests that Sema4D-Sema7A can interact via SEMA domain and form heterodimers, which is concordant with previous studies about formation of multimeric complexes while binding to their functional receptors ^14, 23, 54^.

How heterodimerization affects Sema4D signaling remains unclear, and cannot be easily predicted by structural models, as Sema4D crystal structures use only extracellular regions. One possibility is that compared to homodimerization, heterodimerization exposes certain amino acids in the Sema4D intracellular domain to enzymatic activities. Sema4D has a 107 amino acid cytoplasmic tail where post-translational modifications have been observed, and have been implicated in the cleavage of Sema4D ^55–61^.

Some evidence of receptor functions of Sema4D has also been reported in non-neuronal cell types ^62^, a notable example being in γδ T lymphocytes where antibody-mediated activation of Sema4D resulted in phosphorylation of ERK, dephosphorylation of cofilin and activation of integrins resulting in cell rounding required for wound healing ^63^.

The hSema4D-497P mutation is an exceptional example of the consequence of a single amino acid substitution. While hSema4D-497P can homo- and hetero- dimerize and is predicted to form almost the identical protein structure, it is not correctly trafficked to the membrane and shows evidence of incomplete glycosylation, yielding predominantly the immature (∼120kDa) version of the protein. Moreover, it is worth noting that the 120kDa and 150kDa versions of Sema4D we observe here are both full length (containing the cytoplasmic domain), given we detect both molecular weights when using a C-Terminal flag tag (Figure 4B). Notably, we could never detect a C-terminal cleaved fragment despite our best efforts.

In neurons, glycan interactions are known to be essential in regulating both the correct sorting of proteins to the plasma membrane and for sorting between dendritic and axonal compartments^64^. For example, incomplete glycosylation of the neurotrophin receptor TrkA, results in an immature, low molecular weight protein that prematurely exists the golgi before reaching the *trans*-Golgi network (TGN). When glycosylated, the heavier form of TrkA reaches the TGN and is incorporated into exocytotic vesicles^52^. In another study, N-glycosylation of Neural L1 is absolutely required for its sorting to the axon^65^. It is likely a similar case with the Semaphorin family, however this sorting may also select between secretory (forward) and membrane bound (forward or reverse) signaling. Indeed, it has been reported that Sema5A may function as an attractant or as a repellent, depending on its glycosylation state ^66^. To date, six O-linked glycosylation sites (T657, T663, S666, T670, T716, S722) have been observed on hSema4D. These sites exist between the IG domain and transmembrane region, an area that is outside the bounds of the published crystal structure.

Given hSema4D-497P can still dimerize but is not correctly trafficked, it functions as a potent dominant negative and is likely to inhibit the normal function of hSema4D in clustering GABAergic synapses reported previously ^49, 67^; which may explain the epilepsy observed in our patient. Similarly, Sema7A-deficient cortices show fewer presynaptic puncta ^68^. Based on our observations that Sema7A is a target of Satb2, and Sema4D is preferentially targeted to the plasma membrane in presence of Sema7A, the origin of epileptic symptoms in Satb2- Associated Syndrome (SAS)^12^ may be partially due to incomplete synapse clustering due to Semaphorin 4D-7A complex deficiency.

The presence or absence of Sema7A may also augment situations where Sema4D has been identified to be instrumental in a neuro-immune axis, where theses Semaphorins have known roles. For example, the Sema4D expressed in microglia has recently been observed to be one of the primary mediators for interactions with plexin-containing astrocytes ^69^. Similarly, neurons express higher levels of Sema4D in Alzheimer’s and Huntington’s disease, and its blockade can reduce astrocyte reactivity ^70^. Further work is required to delineate the pathways by which Sema4D-Sema7A reverse signaling directs morphological changes in neuronal and non-neuronal cells. Finally, a better understanding of how glycosylation of individual members of these multimeric membrane complexes may dictate attraction or repulsion has broad implications in both inflammatory neurodegeneration and neurodevelopmental disorders.

## Supporting information

Key Resources Table

## ACKNOWLEDGEMENTS

We would like to thank Dr. Ingo Bormuth for helping with the initial license applications of IUE experiments. We also thank Magda Krejczy and Prof. Dr. Britta Eickholt for the pCAG-FSF-GFP construct. Additionally, we would like to thank Dr. Mateusz Ambrozkiewicz for helpful scientific discussion. We also thank Roman Wunderlich and Ulrike Gunther for excellent technical support.

## AUTHOR CONTRIBUTIONS

Conceptualization, P.B., A.G.N., and V.T.; Investigation, P.B., A.G.N., R.D., D.L., K.T-T., E.K., E.R., T.B., Pr.Ba, J.E. and M.R.; Formal Analysis, and A.G.N., K.T-T., T.B., Visualization, P.B., A.G.N., E.K., and Pr.Ba.; Writing – Original Draft, P.B. and A.G.N; Writing – Review & Editing, P.B., A.G.N., M.R., V.T.; Resources, P.B., A.G.N., K.Y., T.S., R.G., R.P., J.E.; Supervision, V.T.

## STAR METHODS

### Experimental Procedures

For a list of antibodies and plasmids please see the key resources table. Primers used are listed in Data S1.

### Mouse Mutants

All mouse experiments were carried out in compliance with German law approved by the Landesamt für Gesundheit und Soziales (LaGeSo), Berlin. Wild type, *Satb2^fl/wt^ Nex^wt/wt^*, *Satb2^fl/fl^Nex^wt/wt^*, and *Satb2^fl/wt^Nex^Cre/wt^* were used interchangeably as controls. For ease of depiction and consistency, controls have been labeled as *Satb2^fl/wt^Nex^Cre^* in all figures. The day of vaginal plug was considered embryonic day (E) 0.5. Wild type mice used were from a NMRI background.

### *In utero* electroporation (IUE) & Culture of Organotypic Cortical Slices

The procedure of IUE was performed as described before (Saito, 2006). Briefly, DNA plasmid vectors were injected into the lateral ventricle of mouse embryos at embryonic day 14 (E14) and electroporated brains were isolated and fixed at E18 with 4% Paraformaldehyde in PBS. Fixed brains were sectioned in 80μm thick cryosections. Slice culture for live imaging was prepared according to published protocols with slight modifications ^71^. Briefly, 250μm thick cortical slices were sectioned in low melting media-agarose using a MICROM vibratome 24 hours after IUE at E14.

### Cloning

Primers used for in situ probe templates and expression constructs are found in Data S1. Cloned expression constructs were deposited in addgene (plasmids #190643-190657).

Templates for in situ probes were amplified from mouse cortex cDNA using GoTaq polymerase (Promega) and ligated into the pGEM-T vector (Promega). Linearized probe templates were used to generate RNA probes by in vitro transcription using T7 or SP6 polymerase and dNTP DIG labelling mix (Roche).

Mouse Sema7A was amplified from mouse cortex cDNA using ‘naked’ primers in a first round of PCR with Q5-HF polymerase (NEB), whose product was used as a template in a second PCR using primers containing EcoRI and NotI restriction sites. EcoRI-Sema7A-NotI fragment was ligated into pAL2-T vector (Evrogen) yielding pGEMT-Sema7A. Domain deletion mutagenesis was performed on pAL2-T-Sema7A using the Q5 site-directed mutagenesis kit (NEB). For expression, full length Sema7A or domain mutated Sema7A was ligated into EcoRI/NotI digested pCAG-eGFP (addgene# 164092), giving pCAG-Sema7A. After several failed attempts at an N terminal tag, an internal HA tag was designed after reviewing the crystal structure of Sema7A. Using pAL2-T- Sema7A, an HA tag was inserted using the Q5 mutagenesis kit at an exposed loop at amino acid 352, removing only 2 native amino acids to preserve the loop. This HA tagged form was then ligated into EcoRI linearized pCAG-eGFP to yield pCAG-HA-Sema7A.

As above, mouse Sema4D was amplified with Q5 polymerase using ‘naked’ primers, and a second round of PCR added NheI and NotI restriction sites to the amplicon, which was ligated into the pAL2-T vector (Evrogen) giving pAL2-T-NheI-Sema4D-NotI. After observing which amino acid begins the published structure of Sema4D and factoring where cleavage of the signal peptide (SP) is predicted to occur (SignalP-5.0), a myc tag was inserted prior to the phenylalanine at codon 23 using Q5 mutagenesis (NEB). Digested NheI-SP-Myc-Sema4D-NotI was then ligated into NheI/NotI digested pCAG-MCSAN, a pCAG vector with a modified MCS. pCAG-Sema4D-flag was cloned by amplifying from pAL2- T-Sema4D with primers containing flag sequence and also ligated into NheI/NotI digested pCAG-MCSAN.

Human myc- and flag-tagged versions of Sema4D were designed similar to mouse sp-myc-Sema4D, where the tag is inserted after the predicted cleavage point of the signal peptide. hSema4D was amplified in two fragments ‘A’ and ‘B’ from a human cDNA library using GXL polymerase with the tag added via the primer to the first fragment. ‘A’ and ‘B’ fragments were inserted to EcoRI digested pCAG using NEBuilder to give pCAG-hSema4D. Similarly, the hSema4D-Q497P point mutation was introduced by amplifying two fragments from hSema4D and recombined into EcoRI opened pCAG- using NEBuilder (NEB).

### Chromatin Immunoprecipitation & qPCR

#### Preparation of Chromatin

E18 cortex was collected and snap frozen in liquid nitrogen and stored at -80 until use. ∼75mg of tissue was thawed by immediately placing in 1ml of pre-cooled DMEM, and was homogenized by pipetting, and brought to a final volume of 2ml. Tissue was then cross linked by addition of 200μl of freshly made 10X crosslinking solution (16% formaldehyde, methanol-free (Thermo #28906), 1mM EDTA, 0.5mM EGTA, 100mM NaCl, 50mM HEPES-KOH pH 7.5) and rotated for 10 minutes. Fixation was stopped by addition of 250μl 1.25M glycine, rotating 5 minutes. Following a 5 minute centrifugation (1350g, 4°C), tissue was resuspended in 10ml ice-cold lysis buffer 1 (140mM NaCl, 1mM EDTA, 10% Glycerol, 0.5% NP-40, 0.25% Triton X-100, 10mM Sodium Butyrate, 50mM HEPES-KOH pH 7.55 with 1X Protease Inhibitor Cocktail (Roche) and rotated for 10 minutes at 4°C. Following a 5 minute centrifugation (1350g, swing bucket rotor, 4°C), cell pellet was resuspended in 10ml lysis buffer 2 (cell lysis, 10mM Tris-HCl, pH 8.0, 200mM NaCl, 1mM EDTA, 0.5mM EGTA, 10mM Sodium Butyrate with 1X Protease Inhibitor Cocktail) and rotated 10 minutes at 4°C. 10ul was removed to count nuclei in a hemocytometer to confirm optimal concentration (aiming for 500ul lysis buffer 3 for every 10^6^ nuclei). Following a 5 minute centrifugation (1350g, swing bucket rotor, 4°C), nuclei were incubated for 10 minutes on ice in 100ul lysis buffer 3 (10mM Tris-HCl, pH 8.0, 100mM NaCl, 1mM EDTA, 0.5mM EGTA, 0.1% Sodium

Deoxycholate, 0.5% N-lauroyl sarcosine, 10mM sodium Butyrate with 1X Protease inhibitor Cocktail) supplemented with 1% SDS. Prior to shearing, lysate was diluted in 1400ul lysis buffer 3 not containing SDS. Chromatin was sonicated using a Bioruptor Plus (Diagenode) in v-bottomed Eppendorf tubes for 45 cycles using the high energy setting with a cycle of 30 seconds on 30 second off. 1/10 volume of 10% Triton X-100 was then added to chromatin and then spun at 4°C prior to decanting soluble chromatin into a new tube. Chromatin was then stored at -80 for long term storage. Resulting chromatin displayed 200bp-600bp smear of DNA following purification.

#### Chromatin Immunoprecipitation

Protein G Dynabeads (50μl per IP, 25μl pre-clear per IP = 75μl per IP) were washed 3 times in PBS + 0.25% BSA. Chromatin was pre-cleared by incubating with 25μl of washed beads and rotating 2hrs at 4°C. At the same time 12μg Rabbit anti-Satb2 antibody (self-generated, against the peptide QQSQPTKESSPPREEA) was incubated with 50ul of washed beads per IP, rotating 2hrs at 4°C. 10% of pre-cleared chromatin was then set aside as input. Excess buffer was removed from the antibody-bead mixture and incubated with the pre-cleared chromatin rotating overnight at 4°C. Following the IP, beads were washed 6 times in RIPA (10mM Tris-HCl, pH 8.0, 140mM NaCl, 1mM EDTA, 0.5mM

EGTA, 1% Triton X-100, 0.1% Sodium Deoxycholate, 0.1% SDS). Beads were then washed once in TE buffer + 50mM NaCl, and following a 3 minute spin @ 960g, all remaining supernatant was removed. 210ul of elution buffer (50mM Tris-HCl, pH 8.0, 10mM EDTA, 1% SDS) was added to elute antibody-chromatin complexes from the beads at 65°C for 15 minutes. Supernatant was then transferred to a new tube.

#### DNA Clean Up

Eluted chromatin and Input chromatin were processed simultaneously. Chromatin was mixed with 1/10 volume of 5M NaCl and incubated overnight at 65°C. 4ul of RNase A was then added and incubated for 30 minutes at 37°C. 4ul of Proteinase K was then added and incubated for 30 minutes at 55°C. Samples where then transferred to pre-spun phase lock tubes (5Prime, # 2302810) and an equal volume of Phenol:chloroform saturated with TE buffer (OmniPur®, Millipore #6805) was mixed by inversion prior to centrifugation at max speed. The aqueous phase was drawn off into a new epi and 1/10 volume of 3M Sodium Acetate (pH 5.2), 2.5 volumes of 100% EtOH and Glycogen (to a final concentration of 0.05μg/ul) were added. Tubes were vortexed well and DNA was allowed to precipitate overnight at -20°C. DNA pellet was rinsed in ice cold 70% EtOH and dried until clear, then resuspended in 30ul ddH2O.

#### ChIP-qPCR

The following primers were used for 150- 200 bp fragment amplification within the selected region of Sema7A Intron1: F- CAGCCTAGTGTTGGGATGGT, R- ACAAGCAGGCTTGATTCCAT, and for the selected region of Sema7A 5’ Transcription Start Site (TSS): F- CGGGTAGCGAAGGTTTTCCT, R- CAGCCTTTTCTAGCTTTGCCG.

qPCR was performed using Sybr Green qRT-PCR Mastermix reagent. Efficiency of the primers was first tested using cDNA from cortex, measuring 5 serial dilutions (1:5) and analyzed on a StepOne Plus (Applied Biosystems) using the StepOne Software. The qPCR product was also run on a 1,5% Agarose gel to confirm amplicon size. qPCR was performed on 1 μl of 1:5 diluted cDNA from ChIP preparation, and 1:5 dilution of 1% Input.

Enrichment was calculated by first normalizing 1% of input used to 100% using equation (1), replicate 100% input Ct values (CtI) were then averaged to get a mean adjusted input (2). Enrichment was calculated as a percentage of input by calculating the ΔCt of IP’ed values compared to the adjusted input Ct (3).

1. Adjusted Input: ꓯ CtI; (CtI – 6.644) = CtI adjusted
2. Mean adjusted Input = Σ(CtI adjusted)/(n CtI adjusted)
3. Fold Enrichment = 2^((Mean adjusted Input Ct) – (IP Ct))

### Culture of Primary Cortical Neurons

For the analysis of axon specification *in vitro*, primary neuronal cultures were prepared as described before with minor modifications ^72^. For PLA experiments, isolated neurons were nucleofected with the selected DNA plasmid according to manufacturer’s protocols (Mouse Neuron Nucleofector Kit, VPG-1001, and Amaxa 2b Nucleofection system, Lonza). For DIV2 and DIV4 measures of polarity in wildtype and Satb2-/- neurons, plasmid DNA was introduced by chemical transfection using Lipofectamine 2000 (Thermo Fischer) according to the manufacturer’s protocol. Isolated neurons were transfected with the appropriate plasmids as above. *Satb2^fl/wt^* (for simplicity here labelled as WT), *Satb2^fl/fl^* (labelled as Satb2 -/-) and Sema7A rescue neurons were cultured for 2 days and 4 days in vitro (DIV2, DIV4, respectively). Neurons were then cultivated in Neurobasal media (Gibco) supplemented with Glutamax, Penicillin-Streptomycin, and B27at 37°C in 5% CO2. Neurons were fixed at the appropriate time using 4% paraformaldehyde (PFA) for 20 minutes at room temperature. For quantification of number of axons, cells were immunostained with markers for axons and dendrites and counted neurites were positive for Tau1 (1:1000, Millipore) and negative for MAP2 (1:1000, Novus). For unbiased measurements, we used a standardized pixel cross along with pre-defined criteria to measure the acute angle of the centrosome to the closest process at DIV2.

### De-Glycosylation Assay

Two wells of a six well plate of HEK293T cells were transfected with hSema4D or hSema4D-Q497P using lipofectamine 2000 and cultured for 24hrs. Each well was lysed in 100ul Flag lysis buffer (50mM Tris pH 7.4, 150mM NaCl, 1mM EDTA, 1% Triton X100) supplemented with 1X Protease Inhibitor Cocktail (Roche). For each enzymatic condition including undigested, 9ul of lysate was incubated with Glycoprotein Denaturing Buffer (NEB) & heated for 10 minutes at 100°C. Denatured lysates were then chilled on ice & briefly centrifuged before incubating with respective enzymes. For PNGase F lysates, reaction was brought to 20 μl by addition of 2μl Glycobuffer 2 (NEB), 2μl 10% NP-40, 5 μl H2O & 1μl PNGase F (NEB). For O-Glycosidase lysates each reaction was brought to 20ul by addition of 2μl Glycobuffer 2 (NEB), 2μl 10% NP-40, 2μl H2O, 2 μl Neuramidase (NEB) & 2 μl O-Glycosidase (NEB). For Endo H lysates, each reaction was brought to 20μl by addition of 2 μl GlycoBuffer 3 (NEB), 6μl H2O, 2μl Endo H (NEB). Incubation with all three enzymes was performed in Glycobuffer 2. Reactions were incubated for 1hr at 37°C, and supplemented with 7ul of 4X Lammeli Buffer prior to boiling at 95°C for 5 minutes for SDS-PAGE and western blot.

### Immunofluorescence, *In situ* Hybridization (ISH) & Proximity Ligation Assay (PLA)

Immunofluorescence was performed in 2% BSA in PBS containing 1% Triton X-100 (Blocking solution). If Draq5 was to be used for nuclear stain, it was included in the blocking solution with secondary antibodies. *In situ* hybridization was performed according to ^73^. For the Proximity Ligation Assay (PLA), cultured neurons were rinsed once in PBS containing 1 mM MgCl2 and 0.1 mM CaCl2 (PBS-MC) and fixed in PBS-MC containing 4% sucrose and 4% PFA for 20 minutes. PLA was performed according to manufacturer’s instructions (Sigma). Following amplification, coverslips with PLA neurons were incubated in blocking solution for 30 minutes and then incubated with chicken anti-MAP2 (Novus) overnight in blocking solution at 4°C, followed by secondary antibody incubation and final mounting of coverslips for imaging.

### Microscopy & Image Processing

Imaging of fixed cortical sections was performed using an SL-1 confocal microscope (Leica). Specifically, after z stack acquisition, images were flattened to a maximum projection. Neuronal cell culture images and slice culture live imaging were carried out with a Spinning disc microscope (Zeiss).

### Co-Immunoprecipitations & Surface Biotinylation Assay

Twenty-four hours prior to isolation of protein, HEK293T or Hela cells were transfected (Lipofectamine 2000) with the appropriate DNA constructs. pCAG-MCSAN empty vector was used to equilibrate the total amount of DNA to 1μg/μl in all transfections, where 3ug of total DNA was applied to one well of a six well plate.

For lysis, media was removed and cells were lysed in 200μl ice-cold Flag lysis buffer (50mM Tris pH 7.4, 150mM NaCl, 1mM EDTA, 1% Triton X100) supplemented with 1X Protease inhibitor cocktail (Roche). For phosphorylation-sensitive lysates, flag lysis buffer was supplemented to contain final concentrations of 1X Phosstop (Roche), 5μg/ml Pepstatin, 10μg/ml Leupeptin, 2.5mM Sodium-Ortho-Vanadate, 10mM Benzamidine, 1mM β-Glycerophosphate, 5mM NaF, 5μg/ml Aprotinin. Lysates were agitated 4x by gentle pipetting and nuclei were pelleted by 10mins centrifugation at max speed on a tabletop centrifuge at 4°C. Supernatant lysate was then transferred to a new tube and protein concentration was measured by BCA.

For Biotinylation, Hela cells transfected in a six well plate were rinsed once in 0.5ml ice-cold rinsing solution (0.1mM CaCl2, 1mM MgCl2 in PBS) and then incubated for 15 minutes with 0.5ml Sulfo-NHS-SS biotin (1 mg/ml in rinsing solution) on ice. Cells were then rinsed briefly in quenching solution (Rinsing solution + 100mM Glycine pH 2.5) on ice, then rinsed again 10 minutes in quenching solution at 4 degrees to quench unbound biotin. Cells were rinsed once more in rinsing solution and then lysed in Flag lysis buffer supplemented with 1X Protease inhibitor cocktail (Roche), spun 10 minutes (max) at 4°C, then supernatant transferred to a new tube and protein concentration measured by BCA. 50μg of lysate was set aside as whole cell lysate, and 50μg of lysate was incubated with 40μl of avidin agarose beads (NeutrAvidin, Thermo #29201) for 2 hrs at 4°C. Following a brief centrifugation, the supernatant (containing unbiotinylated intracellular fraction) was retained. Beads containing the biotinylated cell surface fraction were then washed 2x with Flag buffer and 2x with TBS, spinning down between washes at 6000rpm for 5 seconds on a tabletop centrifuge. Samples were then boiled at 95°C for 5 minutes in lammeli buffer before running on SDS-PAGE.

### Data Analysis and Statistics

Statistics were performed using either GraphPad Prism 9.0 or using R. Values and statistical details for all experiments are listed in the figure legends. Normality was tested using the Shapiro-Wilk test of normality. Normally distributed data was analyzed using ANOVA with Bonferoni post hoc test for multiple comparisons. Non-normally distributed data was analyzed using Kruskal-Wallis test with Dunn’s Multiple comparison. Probabilities were consistently presented as follows: adjusted p < 0.001 = ***, p <0.01 = **, p < 0.05 = *.

### Structural Modeling

#### Sema4D-Sema7A Heterodimer Superposition

Protein structures of Semaphorin 4D (blue) and Semaphorin 7A (yellow) were superimposed using Discovery Studio 5.0 to investigate the structural similarity. The crystal structures were retrieved from the RCSB Protein Data Bank (PDB) database ^74^, using the accession ID for Semaphorin 4D and Semaphorin 7A - 3OL2 and 3NVQ, respectively. The alignment was based on specified residues. The best superimposition is determined based on a least-squares algorithm minimizing the interatomic distances of the equivalent atoms of the superimposed PDB structures. Only the C^α^ atoms were used since they show the least variation and are well determined ^75^. The superimposition of both the protein structures showed a minor deviation (RMSD:0.64 Å) in the C^α^ atoms of the proteins.

#### Prediction of hSema4D-Q497P structure using Alphafold2

The colabfold implementation of alphafold2 using MMSeqs2 (https://colab.research.google.com/github/sokrypton/ColabFold/blob/mai n/AlphaFold2.ipynb) ^51^ was used to predict the folding of monomers of hSema4D and hSema4D-497P using protein sequences without signal peptides and default parameters. With predicted monomers, the published crystal structure of Sema4D (1OLZ ^76^) was used as a scaffold for dimerization using PyMOL ^77^. Codon 497 and relevant post-translational sites (phosphosite.org, ^78^) were highlighted and images exported.

### Analysis of Published Datasets

Computation has been performed on the HPC for Research/Clinic cluster of the Berlin Institute of Health. Raw read data (fastq) from paired-end RNAseq experiments from P0 wiltdtype (SRR2027113, SRR2027115, SRR2027117) and Satb2 -/- (SRR2027107, SRR2027109, SRR2027111) cortices was downloaded from GEO. Reads were aligned to mm10 using STAR and a count matrix generated using built-in --quantMode GeneCounts. RPKM for the Sema7A gene was calculated using the RPKM function in edgeR. For Satb2 ChIPseq, processed narrowPeak files from GSM2046905 were visualized in IGV browser.

### Exome Sequencing

Trio exome sequencing was performed using a Sure Select Human All Exon 60Mb V6 Kit (Agilent) for enrichment and an Illumina NovaSeq6000 system (Illumina, San Diego, California, USA). Reads were aligned to the UCSC human reference assembly (hg19) with BWA v.0.5.8.1. Single-nucleotide variants (SNVs) and small insertions and deletions were detected with SAMtools v.0.1.7. Copy number variations (CNVs) were detected with ExomeDepth and Pindel. Variant prioritization was performed based on an autosomal recessive (MAF <0.1%) and autosomal dominant (de novo variants, MAF <0.01%) inheritance. This study was approved by the Ethics Committee of the Technical University of Munich, Munich, Germany (#5360/12S).

## SUPPLEMENTARY FIGURES

**Figure S1.**
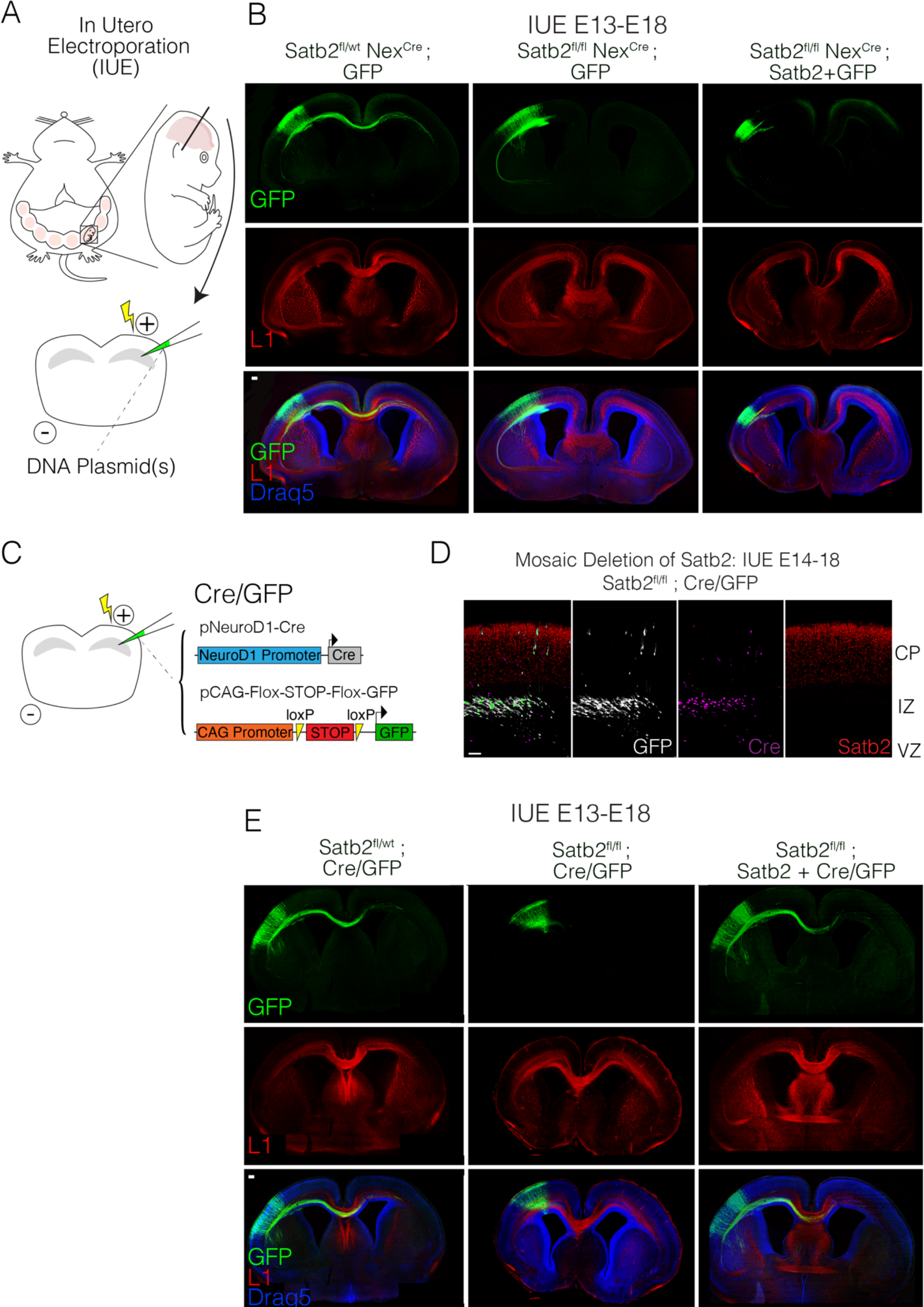
(previous page). *Cell-*intrinsic Satb2 deficits can be rescued by re-expressing Satb2. (A) Schematic showing In Utero Electroporation (IUE) technique involving the introduction of plasmid DNA into the lateral ventricle of live embryos using a capillary and an electric pulse. (B) Satb2 re-expression cannot rescue migration and axon extension in a completely Satb2-deficient cortex. (Left) IUE of GFP into a Satb2^fl/wt^ Nex^Cre^ cortex results in normal migration and axon projection via the corpus callosum. IUE of GFP into a Satb2^fl/fl^ Nex^Cre^ cortex results in cells that project via the internal capsule. Re-expression of Satb2 along with GFP into a Satb2^fl/fl^ Nex^Cre^ cortex abolishes axon projections via the internal capsule and does not rescue axon projection to the midline (Right). L1 marks axonal tracts whereas Draq5 marks nuclei. Scale bar 100μm. (C) Strategy for mosaic deletion of Satb2 from newly born neurons by in utero electroporation. pNeuroD1-Cre confines Cre to post-mitotic neurons expressing NeuroD1. Upon Cre expression, floxed genomic Satb2 is excised, and the STOP cassette from pCAG-FSF-GFP is excised, resulting in GFP expression. (D) Cre/GFP plasmid strategy ensures all green cells observed are Cre positive and also Satb2 negative. These Cre positive, Satb2 negative cells that are labelled with GFP in the Ventricular zone (VZ) do not enter the cortical plate (CP) and remain in the Intermediate Zone (IZ). (E) Satb2 re-expression can rescue cells where Satb2 has been deleted in a mosaic fashion. Cre-injected neurons in *Satb2^fl/wt^* (left) migrate and project axons normally. IUE of Cre into a *Satb2^fl/fl^* cortex (middle) results in halted migration and an absence of axon projection. (Right): when *Satb2* is deleted by Cre and re-introduced into a *Satb2^fl/fl^* cortex, migration into the cortical plate and axonal projections to the midline are restored. Scale bar 100 µm.

**Figure S2.**
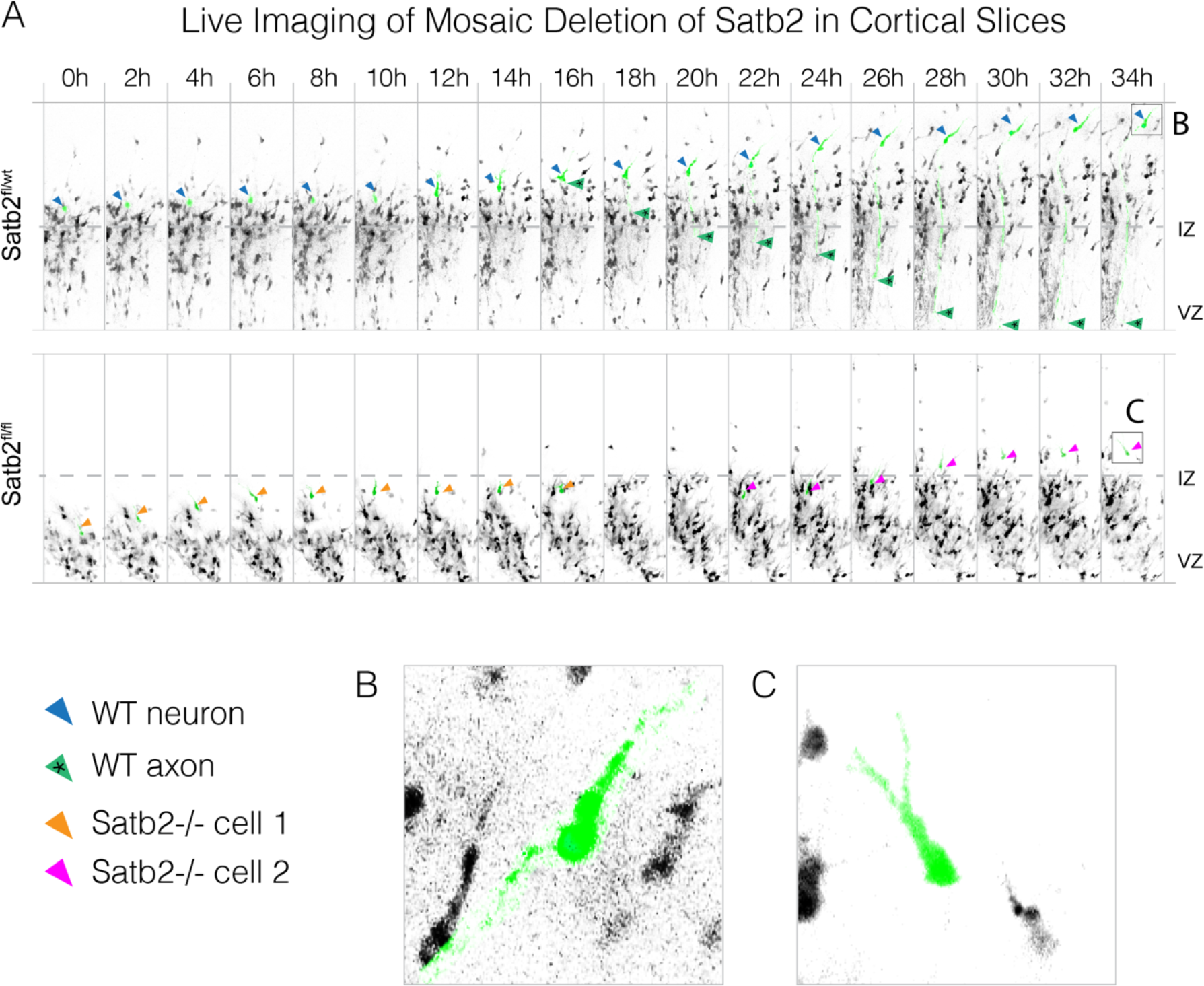
Live imaging of *Satb2* mosaic deletion in organotypic cortical slices. (A) pNeuroD1-Cre + pCAG-FSF-GFP *in utero* electroporation (IUE) visualized in live organotypic slice culture (250 μm thick). A *Satb2^fl/wt^* single cell (pseudocoloured green, yellow arrow) migrates towards the cortical plate and extends an axon (yellow arrow with asterisk) during the imaging time (B). The bottom panel in (A) shows a *Satb2^fl/fl^* pseudocoloured cell (orange arrow) that fails to polarize and remains in the intermediate zone, and another *Satb2^fl/fl^* pseudocoloured cell (magenta arrow) that barely exits the IZ showing a bifurcated leading process (C).

**Figure S3.**
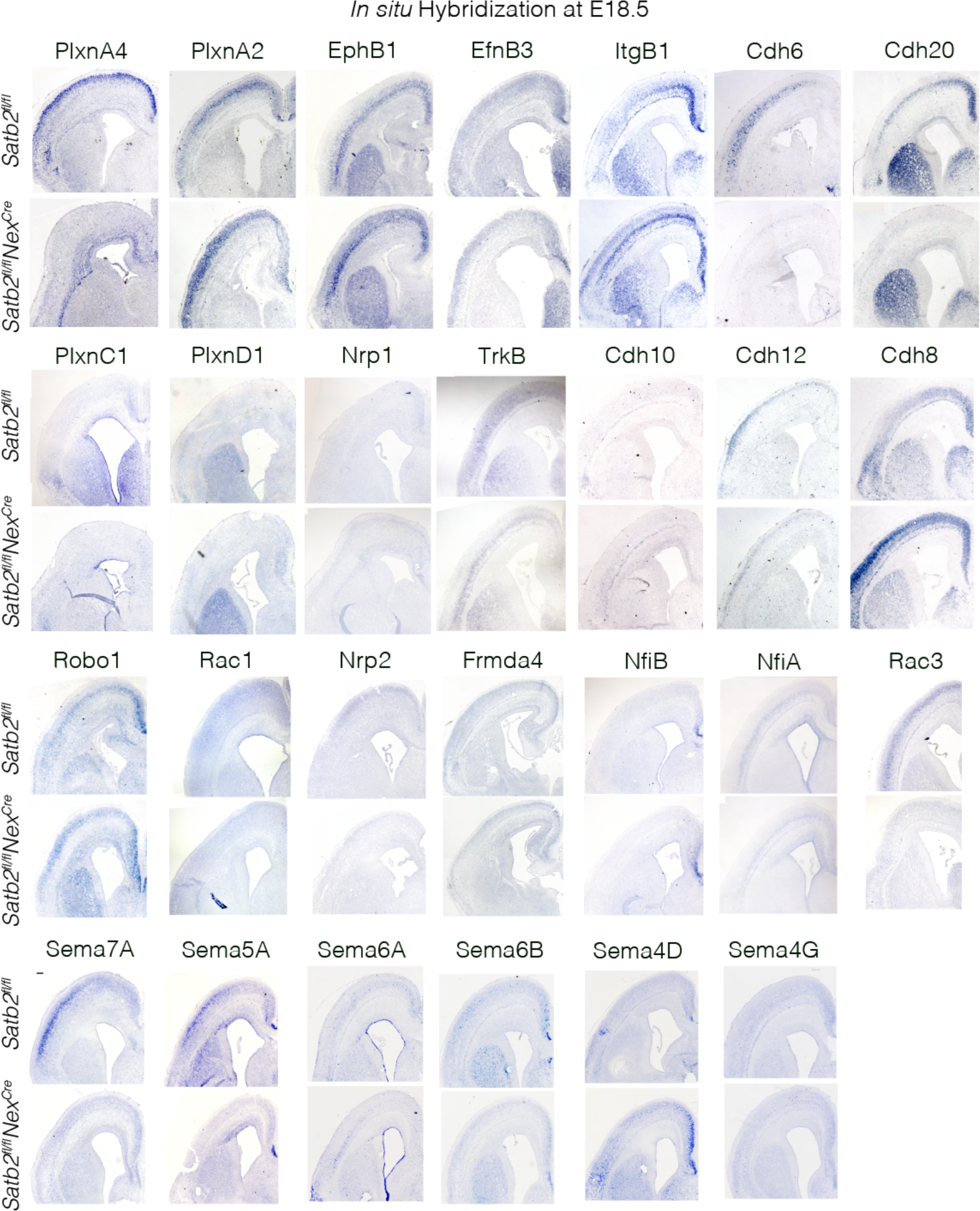
*In situ hybridization* screen aiming to identify *Satb2-* downstream related genes that are expressed in the cortical plate (CP) and are known to influence neuronal polarization and guidance.

**Figure S4.**
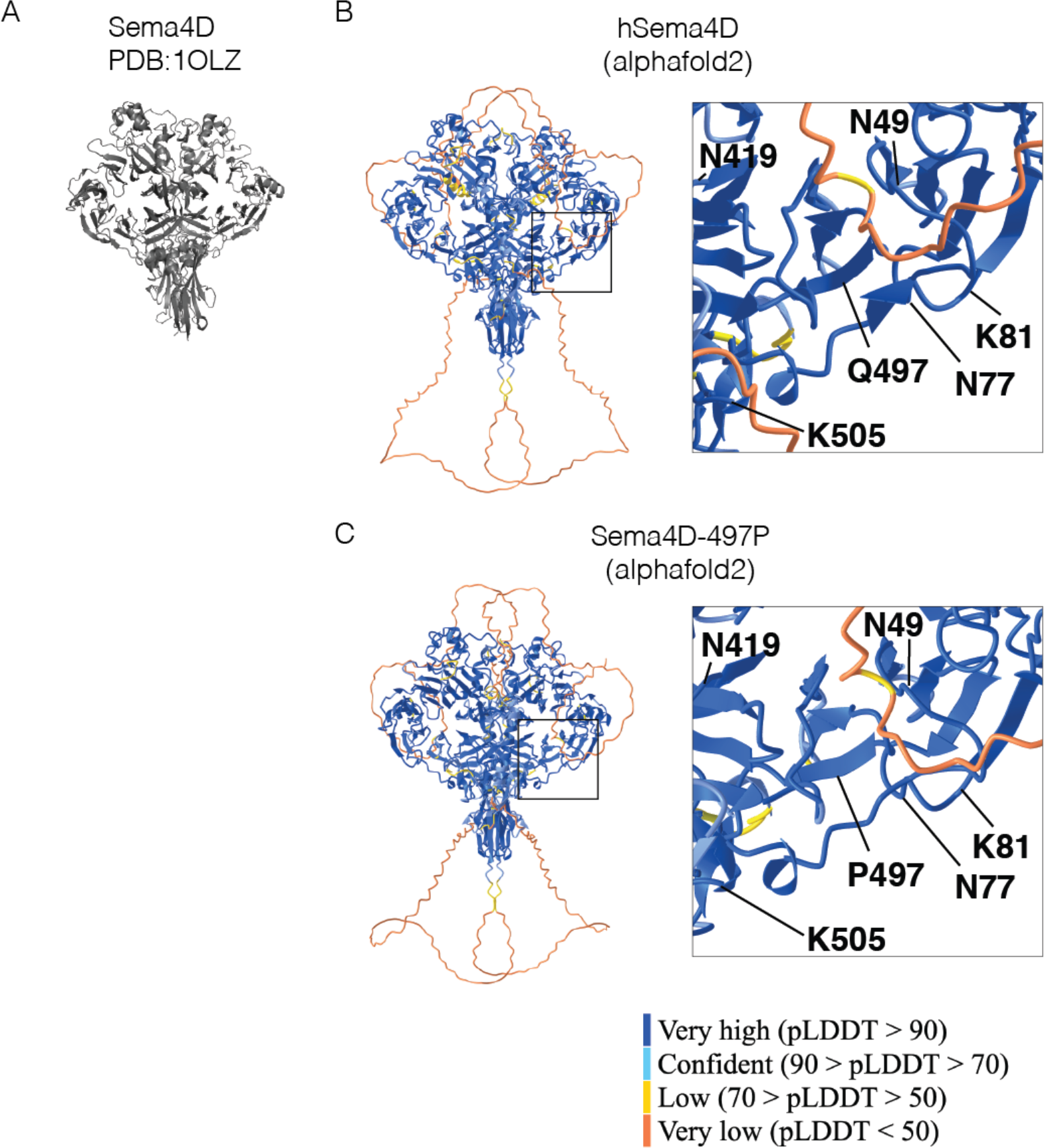
Alphafold2 predicted sema4D structures colored by prediction confidence. (A) Solved crystal structure of human Sema4D. (B) Alphafold2 predicted structure of wild-type human Sema4D. (C) Alphafold2 predicted structure of human Sema4D with 497P mutation.

## Supplemental Note 1

### Case Report: SEMA4D LMU Dr. von Hauner Children Hospital

This boy is the first child of healthy non-consanguineous parents with German- Romanian ancestry. He was born at 40+2 gestational weeks after an uncomplicated pregnancy (weight 3390 g, APGAR 10/10). Delivery was by C-section due to obstructed labor. Newborn hearing and metabolic screening were normal. Due to a mild systolic heart murmur, echocardiography was performed showing a grade 1-2 tricuspid insufficiency. The further postnatal period was unremarkable. At the age of 5 months, the boy presented with a cluster of first generalized tonic-clonic seizures. Electroencephalography (EEG) showed left temporal spikes as well as intermittent slowing on the left parietal side. A brain MRI was performed showing only slightly hyperintense white matter that was interpreted as probably within normal range. Furthermore, he underwent several additional diagnostic procedures (lumbar puncture, laboratory and metabolism tests) that provided normal results. The family history was unremarkable except for the father, who had a simple febrile seizure when he was a baby.

The boy was seizure-free upon anti-seizure treatment with Levetiracetam and Oxcarbazepine. Seizures reoccurred with a monotherapy of Levetiracetam, so monotherapy was switched to Oxcarbazepine. Afterwards he did not show any seizures with Oxcarbazepine monotherapy until the age of 22 months and after that without any medication until the current age of 28 months. Following EEGs showed normal results. Psychomotor development was normal. He was walking at the age of 14 months and spoke short sentences at the age of 2 years. Clinical examination showed no dysmorphic features. Growth was regular, since weight, length, and OFC were 10 kg (17^th^ centile), 80 cm (8^th^ centile), and 51 cm (93^th^ centile) respectively.

Trio-based whole exome sequencing (WES) analysis detected the de novo missense variant (c.1490A>C, p.Gln497Pro) in SEMA4D (NM_006378.3).

## REFERENCES

1. Barnes, A.P., and Polleux, F. (2009). Establishment of Axon-Dendrite Polarity in Developing Neurons. Annual Review of Neuroscience 32, 347–381. 10.1146/annurev.neuro.31.060407.125536.

2. Bradke, F., and Dotti, C.G. (2000). Establishment of neuronal polarity: lessons from cultured hippocampal neurons. Current opinion in neurobiology 10, 574–581.

3. Leone, D.P., Srinivasan, K., Chen, B., Alcamo, E., and McConnell, S.K. (2008). The determination of projection neuron identity in the developing cerebral cortex. Curr Opin Neurobiol 18, 28–35. 10.1016/j.conb.2008.05.006.

4. Leyva-Díaz, E., and López-Bendito, G. (2013). In and out from the cortex: development of major forebrain connections. Neuroscience 254, 26–44. 10.1016/j.neuroscience.2013.08.070.

5. Moldrich, R.X., Gobius, I., Pollak, T., Zhang, J., Ren, T., Brown, L., Mori, S., De Juan Romero, C., Britanova, O., and Tarabykin, V. (2010). Molecular regulation of the developing commissural plate. Journal of Comparative Neurology 518, 3645–3661.

6. Paolino, A., Fenlon, L.R., Suárez, R., and Richards, L.J. (2018). Transcriptional control of long-range cortical projections. Current Opinion in Neurobiology 53, 57–65.

7. Alcamo, E.A., Chirivella, L., Dautzenberg, M., Dobreva, G., Fariñas, I., Grosschedl, R., and McConnell, S.K. (2008). Satb2 Regulates Callosal Projection Neuron Identity in the Developing Cerebral Cortex. Neuron 57, 364–377. 10.1016/j.neuron.2007.12.012.

8. Britanova, O., de Juan Romero, C., Cheung, A., Kwan, K.Y., Schwark, M., Gyorgy, A., Vogel, T., Akopov, S., Mitkovski, M., Agoston, D., et al. (2008). Satb2 Is a Postmitotic Determinant for Upper-Layer Neuron Specification in the Neocortex. Neuron 57, 378– 392. 10.1016/j.neuron.2007.12.028.

9. Döcker, D., Schubach, M., Menzel, M., Munz, M., Spaich, C., Biskup, S., and Bartholdi, D. (2014). Further delineation of the SATB2 phenotype. European Journal of Human Genetics 22, 1034–1039.

10. Zarate, Y.A., and Fish, J.L. (2017). *SATB2* -associated syndrome: Mechanisms, phenotype, and practical recommendations. American Journal of Medical Genetics Part A 173, 327–337. 10.1002/ajmg.a.38022.

11. Zarate, Y.A., Perry, H., Ben-Omran, T., Sellars, E.A., Stein, Q., Almureikhi, M., Simmons, K., Klein, O., Fish, J., Feingold, M., et al. (2015). Further supporting evidence for the SATB2-associated syndrome found through whole exome sequencing. Am. J. Med. Genet. 167, 1026–1032. 10.1002/ajmg.a.36849.

12. Lewis, H., Samanta, D., Örsell, J.-L., Bosanko, K.A., Rowell, A., Jones, M., Dale, R.C., Taravath, S., Hahn, C.D., Krishnakumar, D., et al. (2020). Epilepsy and Electroencephalographic Abnormalities in SATB2-Associated Syndrome. Pediatric Neurology 112, 94–100. 10.1016/j.pediatrneurol.2020.04.006.

13. Srivatsa, S., Parthasarathy, S., Britanova, O., Bormuth, I., Donahoo, A.-L., Ackerman, S.L., Richards, L.J., and Tarabykin, V. (2014). Unc5C and DCC act downstream of Ctip2 and Satb2 and contribute to corpus callosum formation. Nature Communications 5. 10.1038/ncomms4708.

14. Kolodkin, A.L., and Ginty, D.D. (1997). Steering clear of semaphorins: neuropilins sound the retreat. Neuron 19, 1159–1162.

15. Nakamura, F., Kalb, R.G., Strittmatter, S.M., and others (2000). Molecular basis of semaphorin-mediated axon guidance. Journal of neurobiology 44, 219–229.

16. Tamagnone, L., and Comoglio, P.M. (2004). To move or not to move? EMBO reports 5, 356–361. 10.1038/sj.embor.7400114.

17. Pasterkamp, R.J., and Giger, R.J. (2009). Semaphorin function in neural plasticity and disease. Current Opinion in Neurobiology 19, 263–274. 10.1016/j.conb.2009.06.001.

18. Rizzolio, S., and Tamagnone, L. (2007). Semaphorin signals on the road to cancer invasion and metastasis. Cell adhesion & migration 1, 62–68.

19. Yazdani, U., and Terman, J.R. (2006). The semaphorins. Genome biology 7, 211.

20. Jongbloets, B.C., Ramakers, G.M.J., and Pasterkamp, R.J. (2013). Semaphorin7A and its receptors: Pleiotropic regulators of immune cell function, bone homeostasis, and neural development. Seminars in Cell & Developmental Biology 24, 129–138. 10.1016/j.semcdb.2013.01.002.

21. Jongbloets, B.C., Lemstra, S., Schellino, R., Broekhoven, M.H., Parkash, J., Hellemons, A.J.C.G.M., Mao, T., Giacobini, P., van Praag, H., De Marchis, S., et al. (2017). Stage- specific functions of Semaphorin7A during adult hippocampal neurogenesis rely on distinct receptors. Nature Communications 8, 14666. 10.1038/ncomms14666.

22. Negishi, M., Oinuma, I., and Katoh, H. (2005). Plexins: axon guidance and signal transduction. Cellular and Molecular Life Sciences 62, 1363–1371. 10.1007/s00018-005-5018-2.

23. Tamagnone, L., Artigiani, S., Chen, H., He, Z., Ming, G., Song, H., Chedotal, A., Winberg, M.L., Goodman, C.S., Poo, M., et al. (1999). Plexins are a large family of receptors for transmembrane, secreted, and GPI-anchored semaphorins in vertebrates. Cell 99, 71–80.

24. Perez-Branguli, F., Zagar, Y., Shanley, D.K., Graef, I.A., Chédotal, A., and Mitchell, K.J. (2016). Reverse Signaling by Semaphorin-6A Regulates Cellular Aggregation and Neuronal Morphology. PLOS ONE 11, e0158686. 10.1371/journal.pone.0158686.

25. Hernandez-Fleming, M., Rohrbach, E.W., and Bashaw, G.J. (2017). Sema-1a Reverse Signaling Promotes Midline Crossing in Response to Secreted Semaphorins. Cell Reports 18, 174–184. 10.1016/j.celrep.2016.12.027.

26. Kang, S., Nakanishi, Y., Kioi, Y., Okuzaki, D., Kimura, T., Takamatsu, H., Koyama, S., Nojima, S., Nishide, M., Hayama, Y., et al. (2018). Semaphorin 6D reverse signaling controls macrophage lipid metabolism and anti-inflammatory polarization. Nat Immunol 19, 561–570. 10.1038/s41590-018-0108-0.

27. Sun, T., Yang, L., Kaur, H., Pestel, J., Looso, M., Nolte, H., Krasel, C., Heil, D., Krishnan, R.K., Santoni, M.-J., et al. (2017). A reverse signaling pathway downstream of Sema4A controls cell migration via Scrib. The Journal of Cell Biology 216, 199–215. 10.1083/jcb.201602002.

28. Battistini, C., and Tamagnone, L. (2016). Transmembrane semaphorins, forward and reverse signaling: have a look both ways. Cellular and Molecular Life Sciences 73, 1609– 1622. 10.1007/s00018-016-2137-x.

29. Chilton, J.K. (2006). Molecular mechanisms of axon guidance. Developmental Biology 292, 13–24. 10.1016/j.ydbio.2005.12.048.

30. Dudanova, I., and Klein, R. (2013). Integration of guidance cues: parallel signaling and crosstalk. Trends in Neurosciences 36, 295–304. 10.1016/j.tins.2013.01.007.

31. Kim, S.W., and Kim, K.-T. (2020). Expression of Genes Involved in Axon Guidance: How Much Have We Learned? Int J Mol Sci 21, 3566. 10.3390/ijms21103566.

32. Goebbels, S., Bormuth, I., Bode, U., Hermanson, O., Schwab, M.H., and Nave, K.-A. (2006). Genetic targeting of principal neurons in neocortex and hippocampus of NEX-Cre mice. genesis 44, 611–621. 10.1002/dvg.20256.

33. McKenna, W.L., Ortiz-Londono, C.F., Mathew, T.K., Hoang, K., Katzman, S., and Chen, B. (2015). Mutual regulation between Satb2 and Fezf2 promotes subcerebral projection neuron identity in the developing cerebral cortex. PNAS 112, 11702–11707. 10.1073/pnas.1504144112.

34. Hammal, F., de Langen, P., Bergon, A., Lopez, F., and Ballester, B. (2022). ReMap 2022: a database of Human, Mouse, Drosophila and Arabidopsis regulatory regions from an integrative analysis of DNA-binding sequencing experiments. Nucleic Acids Res 50, D316–D325. 10.1093/nar/gkab996.

35. Britanova, O., Alifragis, P., Junek, S., Jones, K., Gruss, P., and Tarabykin, V. (2006). A novel mode of tangential migration of cortical projection neurons. Developmental Biology 298, 299–311. 10.1016/j.ydbio.2006.06.040.

36. Jaitner, C., Reddy, C., Abentung, A., Whittle, N., Rieder, D., Delekate, A., Korte, M., Jain, G., Fischer, A., Sananbenesi, F., et al. (2016). Satb2 determines miRNA expression and long-term memory in the adult central nervous system. eLife 5, e17361. 10.7554/eLife.17361.

37. Stiess, M., and Bradke, F. (2011). Neuronal polarization: The cytoskeleton leads the way. Developmental Neurobiology 71, 430–444. 10.1002/dneu.20849.

38. de Anda, F.C., Pollarolo, G., Da Silva, J.S., Camoletto, P.G., Feiguin, F., and Dotti, C.G. (2005). Centrosome localization determines neuronal polarity. Nature 436, 704–708. 10.1038/nature03811.

39. Sakakibara, A., Sato, T., Ando, R., Noguchi, N., Masaoka, M., and Miyata, T. (2014). Dynamics of Centrosome Translocation and Microtubule Organization in Neocortical Neurons during Distinct Modes of Polarization. Cerebral Cortex 24, 1301–1310. 10.1093/cercor/bhs411.

40. Yoshimura, T., Arimura, N., and Kaibuchi, K. (2006). Signaling Networks in Neuronal Polarization. Journal of Neuroscience 26, 10626–10630. 10.1523/JNEUROSCI.3824-06.2006.

41. Liu, H., Juo, Z.S., Shim, A.H.-R., Focia, P.J., Chen, X., Garcia, K.C., and He, X. (2010). Structural Basis of Semaphorin-Plexin Recognition and Viral Mimicry from Sema7A and A39R Complexes with PlexinC1. Cell 142, 749–761. 10.1016/j.cell.2010.07.040.

42. Kong-Beltran, M., Stamos, J., and Wickramasinghe, D. (2004). The Sema domain of Met is necessary for receptor dimerization and activation. Cancer Cell 6, 75–84. 10.1016/j.ccr.2004.06.013.

43. Marita, M., Wang, Y., Kaliszewski, M.J., Skinner, K.C., Comar, W.D., Shi, X., Dasari, P., Zhang, X., and Smith, A.W. (2015). Class A Plexins Are Organized as Preformed Inactive Dimers on the Cell Surface. Biophysical Journal 109, 1937–1945. 10.1016/j.bpj.2015.04.043.

44. Janssen, B.J., Malinauskas, T., Weir, G.A., Cader, M.Z., Siebold, C., and Jones, E.Y. (2012). Neuropilins lock secreted semaphorins onto plexins in a ternary signaling complex. Nature structural & molecular biology 19, 1293–1299.

45. Pasterkamp, R.J., Peschon, J.J., Spriggs, M.K., and Kolodkin, A.L. (2003). Semaphorin 7A promotes axon outgrowth through integrins and MAPKs. Nature 424, 398–405.

46. Kozlov, G., Perreault, A., Schrag, J.D., Park, M., Cygler, M., Gehring, K., and Ekiel, I. (2004). Insights into function of PSI domains from structure of the Met receptor PSI domain. Biochemical and Biophysical Research Communications 321, 234–240. 10.1016/j.bbrc.2004.06.132.

47. Liu, H., Juo, Z.S., Shim, A.H.-R., Focia, P.J., Chen, X., Garcia, K.C., and He, X. (2010). Structural Basis of Semaphorin-Plexin Recognition and Viral Mimicry from Sema7A and A39R Complexes with PlexinC1. Cell 142, 749–761. 10.1016/j.cell.2010.07.040.

48. Elhabazi, A., Lang, V., Hérold, C., Freeman, G.J., Bensussan, A., Boumsell, L., and Bismuth, G. (1997). The Human Semaphorin-like Leukocyte Cell Surface Molecule CD100 Associates with a Serine Kinase Activity. J. Biol. Chem. 272, 23515–23520. 10.1074/jbc.272.38.23515.

49. Raissi, A.J., Staudenmaier, E.K., David, S., Hu, L., and Paradis, S. (2013). Sema4D localizes to synapses and regulates GABAergic synapse development as a membrane- bound molecule in the mammalian hippocampus. Mol Cell Neurosci 57, 23–32. 10.1016/j.mcn.2013.08.004.

50. Janssen, B.J.C., Robinson, R.A., Pérez-Brangulí, F., Bell, C.H., Mitchell, K.J., Siebold, C., and Jones, E.Y. (2010). Structural basis of semaphorin-plexin signalling. Nature 467, 1118–1122. 10.1038/nature09468.

51. Mirdita, M., Schütze, K., Moriwaki, Y., Heo, L., Ovchinnikov, S., and Steinegger, M. (2022). ColabFold: making protein folding accessible to all. Nat Methods 19, 679–682. 10.1038/s41592-022-01488-1.

52. Vaegter, C.B., Jansen, P., Fjorback, A.W., Glerup, S., Skeldal, S., Kjolby, M., Richner, M., Erdmann, B., Nyengaard, J.R., Tessarollo, L., et al. (2011). Sortilin associates with Trk receptors to enhance anterograde transport and neurotrophin signaling. Nat Neurosci 14, 54–61. 10.1038/nn.2689.

53. Pasterkamp, R.J. (2012). Getting neural circuits into shape with semaphorins. Nature Reviews Neuroscience 13, 605–618. 10.1038/nrn3302.

54. Gherardi, E., Love, C.A., Esnouf, R.M., and Jones, E.Y. (2004). The sema domain. Current opinion in structural biology 14, 669–678.

55. Barberis, D. (2005). p190 Rho-GTPase activating protein associates with plexins and it is required for semaphorin signalling. Journal of Cell Science 118, 4689–4700. 10.1242/jcs.02590.

56. Basile, J.R., Gavard, J., and Gutkind, J.S. (2007). Plexin-B1 Utilizes RhoA and Rho Kinase to Promote the Integrin-dependent Activation of Akt and ERK and Endothelial Cell Motility. Journal of Biological Chemistry 282, 34888–34895. 10.1074/jbc.M705467200.

57. Elhabazi, A., Delaire, S., Bensussan, A., Boumsell, L., and Bismuth, G. (2001). Biological Activity of Soluble CD100. I. The Extracellular Region of CD100 Is Released from the Surface of T Lymphocytes by Regulated Proteolysis. The Journal of Immunology 166, 4341–4347. 10.4049/jimmunol.166.7.4341.

58. Giordano, S., Corso, S., Conrotto, P., Artigiani, S., Gilestro, G., Barberis, D., Tamagnone, L., and Comoglio, P.M. (2002). The Semaphorin 4D receptor controls invasive growth by coupling with Met. Nature Cell Biology 4, 720–724. 10.1038/ncb843.

59. Nishide, M., Nojima, S., Ito, D., Takamatsu, H., Koyama, S., Kang, S., Kimura, T., Morimoto, K., Hosokawa, T., Hayama, Y., et al. (2017). Semaphorin 4D inhibits neutrophil activation and is involved in the pathogenesis of neutrophil-mediated autoimmune vasculitis. Annals of the Rheumatic Diseases 76, 1440–1448. 10.1136/annrheumdis-2016-210706.

60. Swiercz, J.M., Kuner, R., and Offermanns, S. (2004). Plexin-B1/RhoGEF–mediated RhoA activation involves the receptor tyrosine kinase ErbB-2. The Journal of Cell Biology 165, 869–880. 10.1083/jcb.200312094.

61. Swiercz, J.M., Worzfeld, T., and Offermanns, S. (2008). ErbB-2 and Met Reciprocally Regulate Cellular Signaling via Plexin-B1. Journal of Biological Chemistry 283, 1893– 1901. 10.1074/jbc.M706822200.

62. Kuklina, E.M. (2019). Receptor Functions of Semaphorin 4D. Biochemistry Moscow 84, 1021–1027. 10.1134/S0006297919090049.

63. Witherden, D.A., Watanabe, M., Garijo, O., Rieder, S.E., Sarkisyan, G., Cronin, S.J.F., Verdino, P., Wilson, I.A., Kumanogoh, A., Kikutani, H., et al. (2012). The CD100 Receptor Interacts with Its Plexin B2 Ligand to Regulate Epidermal γδ T Cell Function. Immunity 37, 314–325. 10.1016/j.immuni.2012.05.026.

64. Abad-Rodríguez, J., and Díez-Revuelta, N. (2015). Axon glycoprotein routing in nerve polarity, function, and repair. Trends in Biochemical Sciences 40, 385–396. 10.1016/j.tibs.2015.03.015.

65. McFarlane, I., Breen, K.C., Di Giamberardino, L., and Moya, K.L. (2000). Inhibition of N-glycan processing alters axonal transport of synaptic glycoproteins in vivo. Neuroreport 11, 1543–1547.

66. Kantor, D.B., Chivatakarn, O., Peer, K.L., Oster, S.F., Inatani, M., Hansen, M.J., Flanagan, J.G., Yamaguchi, Y., Sretavan, D.W., Giger, R.J., et al. (2004). Semaphorin 5A Is a Bifunctional Axon Guidance Cue Regulated by Heparan and Chondroitin Sulfate Proteoglycans. Neuron 44, 961–975. 10.1016/j.neuron.2004.12.002.

67. Kuzirian, M.S., Moore, A.R., Staudenmaier, E.K., Friedel, R.H., and Paradis, S. (2013). The class 4 semaphorin Sema4D promotes the rapid assembly of GABAergic synapses in rodent hippocampus. J Neurosci 33, 8961–8973. 10.1523/JNEUROSCI.0989-13.2013.

68. Fukunishi, A., Maruyama, T., Zhao, H., Tiwari, M., Kang, S., Kumanogoh, A., and Yamamoto, N. (2011). The action of Semaphorin7A on thalamocortical axon branching: Thalamocortical axon branching by Sema7A. Journal of Neurochemistry 118, 1008– 1015. 10.1111/j.1471-4159.2011.07390.x.

69. Clark, I.C., Gutiérrez-Vázquez, C., Wheeler, M.A., Li, Z., Rothhammer, V., Linnerbauer, M., Sanmarco, L.M., Guo, L., Blain, M., Zandee, S.E.J., et al. (2021). Barcoded viral tracing of single-cell interactions in central nervous system inflammation. Science 372, eabf1230. 10.1126/science.abf1230.

70. Evans, E.E., Mishra, V., Mallow, C., Gersz, E.M., Balch, L., Howell, A., Reilly, C., Smith, E.S., Fisher, T.L., and Zauderer, M. (2022). Semaphorin 4D is upregulated in neurons of diseased brains and triggers astrocyte reactivity. Journal of Neuroinflammation 19, 200. 10.1186/s12974-022-02509-8.

71. Polleux, F., and Ghosh, A. (2002). The Slice Overlay Assay: A Versatile Tool to Study the Influence of Extracellular Signals on Neuronal Development. Science’s STKE 2002, pl9–pl9. 10.1126/stke.2002.136.pl9.

72. Franzoni, E., Booker, S.A., Parthasarathy, S., Rehfeld, F., Grosser, S., Srivatsa, S., Fuchs, H.R., Tarabykin, V., Vida, I., and Wulczyn, F.G. (2015). miR-128 regulates neuronal migration, outgrowth and intrinsic excitability via the intellectual disability gene Phf6. eLife 4, e04263. 10.7554/eLife.04263.

73. Bormuth, I., Yan, K., Yonemasu, T., Gummert, M., Zhang, M., Wichert, S., Grishina, O., Pieper, A., Zhang, W., Goebbels, S., et al. (2013). Neuronal Basic Helix-Loop-Helix Proteins Neurod2/6 Regulate Cortical Commissure Formation before Midline Interactions. Journal of Neuroscience 33, 641–651. 10.1523/JNEUROSCI.0899-12.2013.

74. Burley, S.K., Bhikadiya, C., Bi, C., Bittrich, S., Chen, L., Crichlow, G.V., Christie, C.H., Dalenberg, K., Di Costanzo, L., Duarte, J.M., et al. (2021). RCSB Protein Data Bank: powerful new tools for exploring 3D structures of biological macromolecules for basic and applied research and education in fundamental biology, biomedicine, biotechnology, bioengineering and energy sciences. Nucleic Acids Research 49, D437–D451. 10.1093/nar/gkaa1038.

75. Sumathi, K., Ananthalakshmi, P., Roshan, M.N.A.M., and Sekar, K. (2006). 3dSS: 3D structural superposition. Nucleic Acids Res 34, W128–132. 10.1093/nar/gkl036.

76. Love, C.A., Harlos, K., Mavaddat, N., Davis, S.J., Stuart, D.I., Jones, E.Y., and Esnouf, R.M. (2003). The ligand-binding face of the semaphorins revealed by the high-resolution crystal structure of SEMA4D. Nat Struct Biol 10, 843–848. 10.1038/nsb977.

77. Schrödinger, LLC (2015). The PyMOL Molecular Graphics System, Version 1.8.

78. Hornbeck, P.V., Zhang, B., Murray, B., Kornhauser, J.M., Latham, V., and Skrzypek, E. (2015). PhosphoSitePlus, 2014: mutations, PTMs and recalibrations. Nucleic acids research 43, D512–D520.

