## Supplementary material for "Sema7A and Sema4D Heterodimerization is Essential for Membrane Targeting and Neocortical Wiring": Key Resources Table

| REAGENT or RESOURCE | SOURCE | IDENTIFIER |
| --- | --- | --- |
| Antibodies | | |
| Rabbit anti-Satb2 | Tarabykin Laboratory |  |
| Goat anti-GFP | Rockland | 600-101-215M; RRID:[AB_2612804](http://antibodyregistry.org/AB_2612804" \t "_blank) |
| Chicken anti-GFP | Abcam | ab13970 |
| Mouse anti-Myc (9B11) | Cell Signalling | 2276 |
| Rabbit anti-Myc (71D10) | Cell Signalling | 2278 |
| Rabbit anti-HA (C29F4) | Cell Signalling | 3724 |
| Mouse anti-Flag | Cell Signalling | 14793 |
| Rabbit anti-Cdk5rap2 | Kaindl Laboratory |  |
| Rabbit anti-vinculin | Cell Signalling | 4650 |
| Rabbit anti-Beta Catenin (D10A8) | Cell Signalling | 8480 |
| Mouse anti-GM130 | BD Biosciences | 610822 |
| Mouse anti-Cre Recombinase (clone 2D8) | Millipore | MAB3120 |
| Chicken anti-MAP2 | Novus | NB300-213; RRID:[AB_350528](http://antibodyregistry.org/AB_350528" \t "_blank) |
| Mouse anti-TAU-1 (clone PC1C6) | Millipore | MAB3420 |
| Donkey anti-goat AlexaFluor-488 | Jackson Immunoresearch | 705-546-147; RRID:[AB_2340430](http://antibodyregistry.org/AB_2340430" \t "_blank) |
| Donkey anti-goat AlexaFluor-Cy3 | Jackson Immunoresearch | 705-165-147; RRID:[AB_2307351](http://antibodyregistry.org/AB_2307351" \t "_blank) |
| Donkey anti-rabbit AlexaFluor-488 | Jackson Immunoresearch | 711-545-152; RRID:[AB_2313584](http://antibodyregistry.org/AB_2313584" \t "_blank) |
| Donkey anti-rabbit AlexaFluor-Cy3 | Jackson Immunoresearch | 711-167-003; RRID:[AB_2340606](http://antibodyregistry.org/AB_2340606" \t "_blank) |
| Donkey anti-rabbit AlexaFluor-Cy5 | Jackson Immunoresearch | 711-165-152; RRID:[AB_2307443](http://antibodyregistry.org/AB_2307443" \t "_blank) |
| Donkey anti-rat AlexaFluor-Cy3 | Jackson Immunoresearch | 712-165-153; RRID:[AB_2340667](http://antibodyregistry.org/AB_2340667" \t "_blank) |
| Donkey anti-rat AlexaFluor-Cy5 | Jackson Immunoresearch | 712-175-153; RRID:[AB_2340672](http://antibodyregistry.org/AB_2340672" \t "_blank) |
| Donkey anti-mouse AlexaFluor-Cy3 | Jackson Immunoresearch | 715-165-150; RRID:[AB_2340813](http://antibodyregistry.org/AB_2340813" \t "_blank) |
| Donkey anti-mouse AlexaFluor-Cy5 | Jackson Immunoresearch | 715-175-151; RRID:[AB_2340820](http://antibodyregistry.org/AB_2340820" \t "_blank) |
| cDNA | | |
| cDNA clone Sema5A | Source Bioscience | BC065137 |
| cDNA clone Sema6A | Source Bioscience | BC059238 |
| Bacterial and virus strains | | |
| Escherichia coli JM109 competent cells | Promega | L2005 |
| Critical commercial assays | | |
| Duolink^®^ In Situ Red Starter Kit Mouse/Rabbit | Sigma | DUO92101 |
| NEBNext® ChIP-seq Library Prep Reagent set for Illumina | New England Biolabs | E6200 |
| Deposited data | | |
| Satb2 KO RNAseq data | (McKenna et al 2014) | [GSE68911](https://https.ncbi.nlm.nih.gov/geo/query/acc.cgi?acc=GSE68911) |
| Experimental models: Cell lines | | |
| HEK293T | Leibniz Institute DSMZ-German Collection of Microorganisms and Cell Cultures | ACC 635; RRID: CVCL_0063 |
| N2A | Thermo Fischer | RRID: CVCL_0470 |
| Experimental models: Organisms/strains | | |
| Mouse: *Satb2^fl/fl^* (SATB2F) *B6.-Satb2^fl/fl^* | Grossschadel Laboratory | N/A |
| Oligonucleotides | | |
| Information regarding the oligonucleotides used in this study can be found in Data S1 | | |
| Recombinant DNA | | |
| pCAG-HA-Sema7A | This paper | Addgene # 190644 |
| pCAG-Sema7A | This paper | Addgene # 190643 |
| pCAGIG-Sema7A-ΔSEMA | This paper | Addgene # 190647 |
| pCAGIG-Sema7A-ΔPSI | This paper | Addgene # 190649 |
| pCAGIG-Sema7A-ΔIG | This paper | Addgene # 190648 |
| pCAGIG-Sema7A-ΔMEM | This paper | Addgene # 190650 |
| pCAGIG-Sema7A-RGD-KCE | This paper | Addgene # 190651 |
| pCMV-myc-Sema4D | (Raissi et al., 2013) | Addgene # 51599 |
| pCAG-sp-myc-Sema4D | This paper | Addgene # 190645 |
| pCAG-Sema4D-flag (C term) | This paper | Addgene # 190646 |
| pCAG-myc-hSema4D | This paper | Addgene # 190654 |
| pCAG-flag-hSema4D | This paper | Addgene # 190656 |
| pCAG-myc-hSema4D-Q497P | This paper | Addgene # 190655 |
| pCAG-flag-hSema4D-Q497P | This paper | Addgene # 190657 |
| pCAG-FSF-EGFP | Tarabykin Laboratory |  |
| pSuper Neo Gfp-sh-Scramble | (Ambrozkiewicz et al., 2018) |  |
| pLKO-shSema7A | Sigma-Aldrich | TRCN0000067538 |
| pLKO-shSema4D | Sigma-Aldrich | TRCN0000067495 |
| pGEM-T | Promega | A3600 |
| pNeuroD1-Cre | Tarabykin Laboratory |  |
| pCAG-EGFP | (Matsuda and Cepko, 2004) | Addgene #11150 |
| Software and algorithms | | |
| Snakemake | (Köster and Rahmann, 2012) | <https://snakemake.readthedocs.io/en/stable/> |
| STAR | (Dobin et al., 2013) | https://github.com/alexdobin/STAR |
| Bowtie | (Langmead and Salzberg, 2012) |  |
| Deeptools | (Ramírez et al., 2016) | https://deeptools.readthedocs.io/en/develop/ |
| BWA v.0.5.8.1 | (Li and Durbin, 2009) |  |
| Samtools v.0.1.7 | (Li et al., 2009) | Htslib.org |
| ExomeDepth | (Plagnol et al., 2012) | https://github.com/vplagnol/ExomeDepth |
| Pindel | (Ye et al., 2009) | https://github.com/genome/pindel |
| BIOVIA Discovery Studio 5.0 | Dassault Systèmes | https://www.3ds.com/ |
| Google Colab Implementation of AlphaFold2 | DeepMind  & Mirdita et al 2022 | https://colab.research.google.com/github/sokrypton/ColabFold/blob/main/AlphaFold2.ipynb |
| Image Lab 6 | BioRad | https://www.bio-rad.com/ |
| Adobe Photoshop | Adobe | https://www.adobe.com/ |
| Adobe Illustrator | Adobe | https://www.adobe.com/ |
| Fiji/ImageJ | (Schindelin et al., 2012) | <https://imagej.nih.gov/ij/> |
| Other | | |
